# Specific nanoscale synaptic reshuffling and control of short-term plasticity following NMDAR- and P2XR-dependent Long-Term Depression

**DOI:** 10.1101/759191

**Authors:** Benjamin Compans, Magalie Martineau, Remco V. Klaassen, Thomas M. Bartol, Corey Butler, Adel Kechkar, David Perrais, Terrence J. Sejnowski, Jean-Baptiste Sibarita, August B. Smit, Daniel Choquet, Eric Hosy

## Abstract

Long-Term Potentiation (LTP) and Long-Term Depression (LTD) of excitatory synaptic transmission are considered as cellular basis of learning and memory. These two forms of synaptic plasticity have been mainly attributed to global changes in the number of synaptic AMPA-type glutamate receptor (AMPAR) through a regulation of the diffusion/trapping balance at the PSD, exocytosis and endocytosis. While the precise molecular mechanisms at the base of LTP have been intensively investigated, the ones involved in LTD remains elusive. Here we combined super-resolution imaging technique, electrophysiology and modeling to describe the various modifications of AMPAR nanoscale organization and their effect on synaptic transmission in response to two different LTD protocols, based on the activation of either NMDA receptors or P2X receptors. While both type of LTD are associated with a decrease in synaptic AMPAR clustering, only NMDAR-dependent LTD is associated with a reorganization of PSD-95 at the nanoscale. This change increases the pool of diffusive AMPAR improving synaptic short-term facilitation through a post-synaptic mechanism. These results demonstrate that specific dynamic reorganization of synapses at the nanoscale during specific LTD paradigm allows to improve the responsiveness of depressed synapses.

## Introduction

Changes in synaptic efficacy, either by strengthening through long-term potentiation (LTP) or weakening through long-term depression (LTD), are believed to underlie learning and memory. These synaptic plasticity mechanisms rely on the regulation of AMPA-type glutamate receptor number at synapses. In addition to endo- and exocytosis which control the total amount of receptors at the neuronal surface, the equilibrium at the membrane between freely-diffusive and immobilized AMPAR at the PSD regulates synaptic responses (1–5). This equilibrium is under the control of a tightly regulated trapping/untrapping mechanism involving the AMPAR associated-proteins, the post-synaptic scaffolding proteins (e.g. PSD-95) which anchor AMPAR complexes at the PSD, and the balanced activity of phosphatases and kinases regulating the affinity of the trapping (6–8). We and others have demonstrated that synaptic strength is not solely dependent on the quantity of glutamate per pre-synaptic vesicle and the number of post-synaptic AMPARs, but also rely on their nanoscale clusterization with respect to the pre-synaptic active zone (9–12). Modeling sets that long term modifications in synaptic strength as observed with LTD or LTP can results from (i) a change in AMPAR density at synapse, (ii) a variation in receptor amount per cluster or (iii) a modification of the alignment between pre-synaptic release sites and AMPAR clusters (9,13).

This dynamic equilibrium between mobile and trapped AMPARs regulates synaptic transmission properties at multiple timescales. Indeed, we established the role of mobile AMPAR in short term plasticity. This plasticity observed when synapses are stimulated at tens of Hz was previously thought to rely on both presynaptic mechanisms, and AMPAR desensitization properties (14–17). We demonstrated that after glutamate release, desensitized AMPARs can rapidly exchange by lateral diffusion with naïve receptors, increasing the number of activatable receptors for a sequential glutamate release (14,18–20). Systematically, a decrease in the pool of mobile AMPARs using receptor crosslinking (14,18), CaMKII activation (19) or chimeric receptor (18,21) enhances short-term depression. In addition to its role on short-term plasticity, AMPAR lateral diffusion was shown to play a key role during LTP (19,22). We recently demonstrated that altering the synaptic recruitment of AMPARs by interfering with AMPAR lateral diffusion impairs the early LTP (22).

These studies demonstrated that nanoscale regulation of AMPAR organization and dynamic tune synaptic responses. However, few studies have yet addressed the molecular reshuffling induced by LTD. Moreover, LTD is a generic term indicating a decrease in synaptic strength but can be induced by different pathways. To investigate how various LTD protocols are associated with the dynamic reorganization of synaptic AMPAR, we combined live and fixed super-resolution microscopy, a fluorescent glutamate sensor and electrophysiology together with modeling. We tested two well characterized LTD protocols based on the activation of either the NMDARs, by NMDA, or the P2X receptors (P2XRs), by ATP (23–25). We observed that NMDAR-, but not P2XR-dependent LTD triggers specific changes in PSD-95 nanoscale organization and increases AMPAR lateral diffusion. Finally, the latter improves the capacity of depressed synapses to integrate high frequency stimulations.

## Results

### ATP and NMDA treatments both induce a long-lasting decrease in synaptic AMPAR content and miniature amplitude

We performed direct Stochastic Optical Reconstruction Microscopy (dSTORM) experiments and electrophysiological recordings to monitor AMPAR organization and currents following the application of either ATP or NMDA, two well-established chemical protocols to trigger a long lasting synaptic depression (23,24). The fluorescent emission property of individual AMPAR was extracted from isolated receptors present at the membrane surface (11) and used to estimate the density and number of AMPARs in different neuronal compartments. NMDA treatment (30µM, 3min) led to a rapid and stable 25percent decrease in AMPAR density both at the dendritic shaft and into spines (Figure 1S1A-B). As previously described, approximately 50percent of synaptic AMPAR are organized in nanodomains facing pre-synaptic release sites (10,11). NMDAR-dependent LTD was also associated with a rapid (within the first 10 minutes following NMDA application) and stable (up to 3 hours) depletion in AMPAR content per nanodomain (estimated number of AMPAR per nanodomain: t0: 19.77 +/-1.20, t10: 12.82 +/-1.05, t30: 12.59 +/-0.82) (Figure 1A-B). In contrast, the overall nanodomain diameter was preserved as shown by the stability of their full width half maximum (t0: 79.13 +/-1.82 nm, t10: 82.04 +/-2.53 nm, t30: 84.35 +/-2.43 nm) (Figure 1C). This reorganization of synaptic AMPAR was associated with a depression in AMPAR-mediated miniature excitatory post-synaptic current amplitude (mEPSC; Figure 1D-E, t0: 11.34 +/-0.50 pA, t10: 7.60 +/-0.49 pA, t30: 8.00 +/-0.65 pA), which lasted up to 3 hours after NMDA treatment (Figure 1F, t0: 10.98 +/-0.73 pA, t180: 7.06 +/-0.49 pA).

**Fig. 1.**
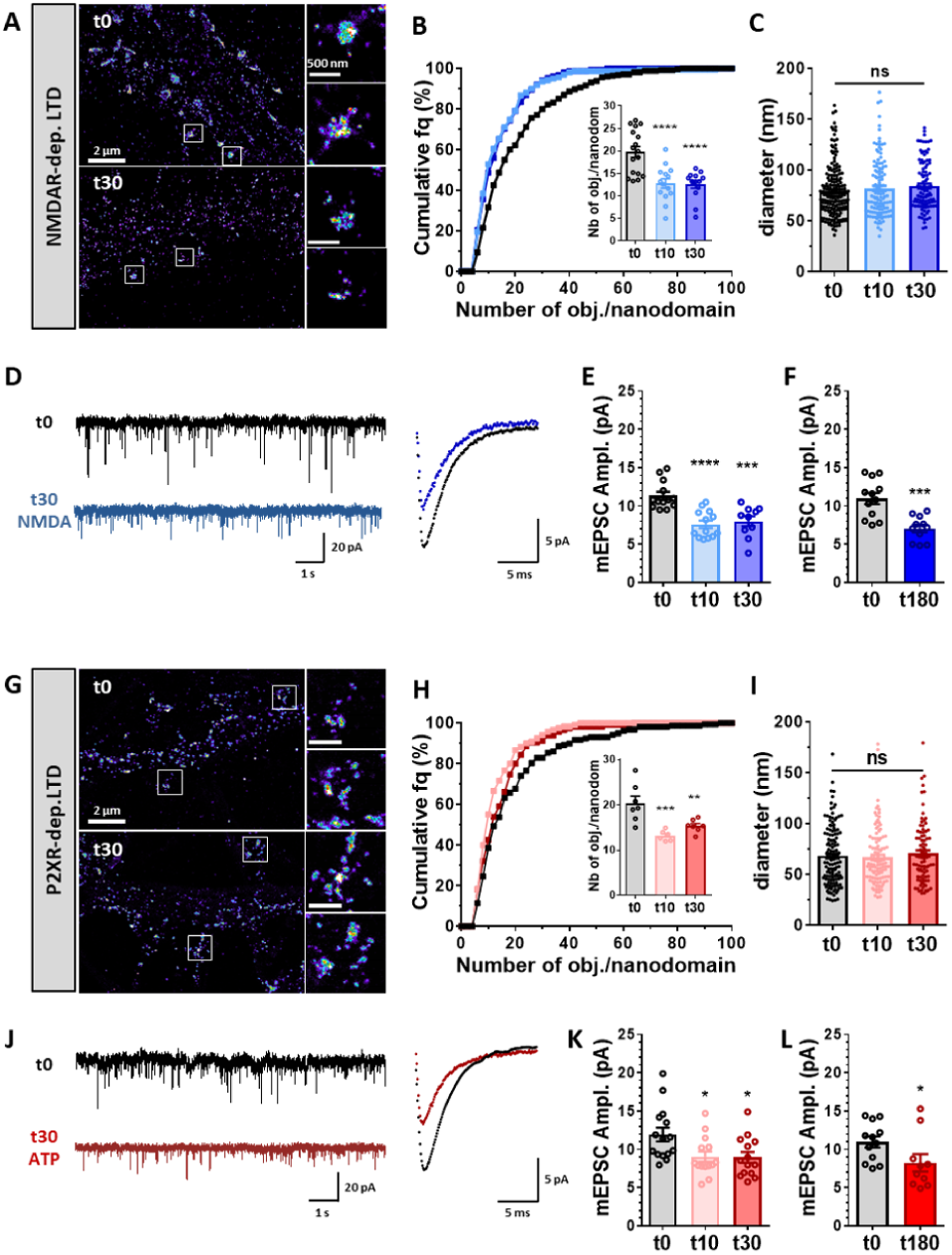
NMDA and ATP application triggers a rapid and long lasting nanoscale re-organization of AMPAR at synapses associated to a long-term synaptic current depression.(A) Example of super-resolution intensity images of a piece of dendrite obtained using dSTORM technique on live stained neurons for endogenous GluA2 containing AMPARs at basal state (t0) or 30 minutes (t30) following NMDA application (30 µM, 3 min). Enlarged synapses are shown on the right. (B) Cumulative distribution of nanodomain AMPAR content (n=275, 159 and 152 for t0, t10 and t30 respectively), and in the inset, the mean per cell. The number of AMPARs per nanodomain was estimated 0, 10 and 30 minutes following NMDA treatment as explained in Nair et al. 2013 (mean +/- SEM, n=17, 14 and 14 respectively, one-way ANOVA, p<0.0001 and Dunnett’s post-test found significant differences between t10 or t30 and t0, p<0.0001). Nanodomain content is significantly decreased 10 and 30 minutes following NMDA treatment compared to non-treated cells. (C) Diameter of AMPAR synaptic nanodomains. Nanodomain sizes were measured by anisotropic Gaussian fitting of pre-segmented clusters obtained on dSTORM images. Nanodomain diameter (mean +/- SEM) 0, 10 and 30 minutes following NMDA treatment are plotted (n=191, 127 and 100 respectively, one-way ANOVA, p=0.2487). Nanodomain size is not affected by NMDA application. (D) Left panel: example of miniature EPSC traces recorded on cultured neurons in basal condition (dark trace) or 30 minutes after NMDA treatment (blue trace). Right panel: Superposition of a mean trace of AMPAR mEPSC in basal (dark) and 30 minutes post-NMDA treatment (blue). (E and F) Average of the mESPC amplitude recorded on neurons 0, 10 or 30 minutes (E) and 180 minutes (F) after NMDA treatment. Miniature EPSC amplitudes are significantly depressed 10 and 30 minutes after NMDA treatment (E, n=13, 13 and 10 respectively, one-way ANOVA p<0.0001 and Dunnett’s post-test found significant differences p<0.0001 and p=0.0003 between t0 and t10, and t0 and t30 respectively), and this depression stays for at least 3 hours (F, n=12 and 11 respectively, t-test p=0.0003). (G) Example of super-resolution intensity images of a piece of dendrite obtained using dSTORM technique on neurons live stained for endogenous GluA2 containing AMPARs at basal state (t0) or 30 minutes (t30) following ATP (100 µM, 1 min). Enlarged synapses are shown on the right. (H) Cumulative distribution of nanodomain AMPAR content (n=158, 120 and 115 for t0, t10 and t30 respectively), and in the inset, the mean per cell. The number of AMPARs per nanodomains was estimated 0, 10 and 30 minutes following ATP treatment (mean +/- SEM, n=7, 6 and 7 respectively, one-way ANOVA, p=0.0006 and Dunnett’s post-test found significant differences between t10 or t30 and t0, p=0.0004 and p=0.0063 respectively). Nanodomain content is decreased 10 and 30 minutes following ATP treatment compared to non-treated cells. (I) Measure of nanodomain diameter is not affected 0, 10 and 30 minutes following ATP treatment (n=130, 112 and 91 respectively, one-way ANOVA, p=0.6391). (J) Left panel: example of miniature EPSC traces recorded on cultured neurons in basal condition (dark trace) or 30 minutes after ATP treatment (red trace). Right panel: Superposition of a mean trace of AMPAR mEPSC in basal (dark) and 30 minutes post-ATP treatment (red). (K and L) Average of the mESPC amplitudes recorded on neurons 0, 10 or 30 minutes (K) and 180 minutes (L) after ATP treatment (100µM, 1min). Synaptic transmission (mEPSCs) is significantly depressed 10 and 30 minutes after ATP treatment (K, n=15, 14 and 14 respectively, one-way ANOVA p=0.0124 and Dunnett’s post-test found significant differences p<0.0214 and p=0.0172 between t0 and t10, and t0 and t30 respectively), and this depression stays for at least 3 hours (L, n=12 and 10 respectively, t-test p=0.0485).

In parallel, we measured the effect of ATP treatment, reported as able to induce a solid and long lasting LTD, on both AMPAR nanoscale organization and mEPSCs. Purinergic receptors from the P2X family were activated using ATP (100µM, 1min) in the presence of CGS15943 (3µM) to avoid adenosine receptor activation (24,26). As for NMDA, ATP treatment triggered a rapid and long-lasting decrease in AMPAR content both at dendritic shafts, spines (Figure 1S1E-F) and nanodomains (estimated number of AMPAR per nanodomain: t0: 20.36 +/-1.61, t10: 13.21 +/-0.46, t30: 15.38 +/-0.53), without affecting their overall organization (t0: 68.19 +/-2.55 nm, t10: 66.81 +/-2.57 nm, t30: 70.62 +/-3.11 nm) (Figure 1G-I). In parallel, this depletion of AMPAR nanodomains was associated with a stable 25percent decrease in mEPSC amplitude (t0: 11.93 +/-0.89 pA, t10: 9.05 +/-0.68 pA, t30: 8.95 +/-0.70 pA) (Figure 1J-L). Altogether, these results indicate that both NMDA- and ATP-induced synaptic depression are associated with a reduction in the number of surface AMPARs at the synapse and on the dendrite, notably leading to a depletion in nanodomain content, without a change in their overall dimensions.

To determine whether the mEPSC amplitude decrease was associated with modifications in current kinetics, which might be caused by changes in the composition of AMPAR complexes, we analyzed both rise and decay times of mEPSCs. We observed no modifications in current kinetics when LTD was induced by either NMDA or ATP (Figure1S2). Interestingly, we observed a transient decrease in the mEPSC frequency after NMDA treatment, which might be explained by the decrease in the number of nanodomains per spine (Figure 1S1C and 1S2). This observation suggests the complete disappearance of some domains. This effect on the nanodomains was not observed when LTD was induced by ATP (Figure1S1G and 1S2). As demonstrated in (9), a decrease in synaptic response could be due to a change in pre- to post-synaptic alignment. To estimate whether NMDA treatment affected the trans-synaptic organization, we performed dual color d-STORM experiments to measure the alignment of the pre-synaptic protein RIM1/2 and the post-synaptic AMPAR. We calculated the centroid to centroid distances between RIM1/2 and GluA2-containing AMPAR clusters at t0, and 10 and 30 minutes after NMDA treatment (Figure 1S3). NMDA treatment did not change significantly the RIM/AMPAR co-organization.

### AMPAR lateral diffusion is increased during NMDAR-dependent LTD but not during P2XR-dependent LTD

It has been reported that both endocytosis and exocytosis are involved in synaptic plasticities and that lateral diffusion is the way for AMPAR to travel from synapses to endo/exocytosis pits. So we next investigated the effect of the two LTD-inducing protocols on AMPAR lateral diffusion. Using the single-particle tracking technique uPAINT (27), we measured endogenous GluA2-containing AMPAR mobility at the cell surface. We acquired time lapse series consisting of 1.5 minute acquisitions performed every 5 minutes for 30 minutes on cells in the basal condition and upon NMDAR- or P2XR-dependent LTD induction. As previously described (11) and illustrated figure 2B and G (control in black line), distribution of AMPAR diffusion coefficients reveal two main populations centered approximatively at 0.8*10-2 µm^2^.s-1 (termed immobile trapped receptors) and 10-1 µm^2^.s-1 (termed mobile receptors). 30 minutes following NMDA treatment, we observed a 35 percent increase in the AMPAR mobile fraction (D coef > 0.02µm^2^.s-1) (t0: 30.11+/-1.69 percent, t30: 40.65+/-2.94 percent) (Figure 2A-C). In contrast, we observed no change in AMPAR lateral diffusion with vehicle (H2O) treatment (t0: 31.17+/- 1.94 percent, t30: 27.71+/- 2.98 percent) (Figure 2D), or 30 minutes after an ATP treatment (t0: 31.19+/- 2.58 percent, t30: 30.44 +/- 1.40 percent) (Figure 2F-J). Similar results were obtained for synaptic trajectories of the GluA2-containing AMPAR (Figure 2S1). This increase in AMPAR mobility following NMDAR-dependent LTD induction takes place progressively along the first 20 minutes following NMDA treatment (Figure 2D) and remains stable up to 3 hours (Figure 2E; t0: 27.25 +/- 3.41 percent, t180: 38.57 +/- 2.56 percent). On the contrary, neither control nor ATP treatment induced such changes in AMPAR mobility (Figure 2I-J). Moreover, in the presence of APV (50 µM), a specific NMDAR antagonist, no increase in AMPAR mobility was observed after NMDA treatment (Figure 2S2).

**Fig. 2.**
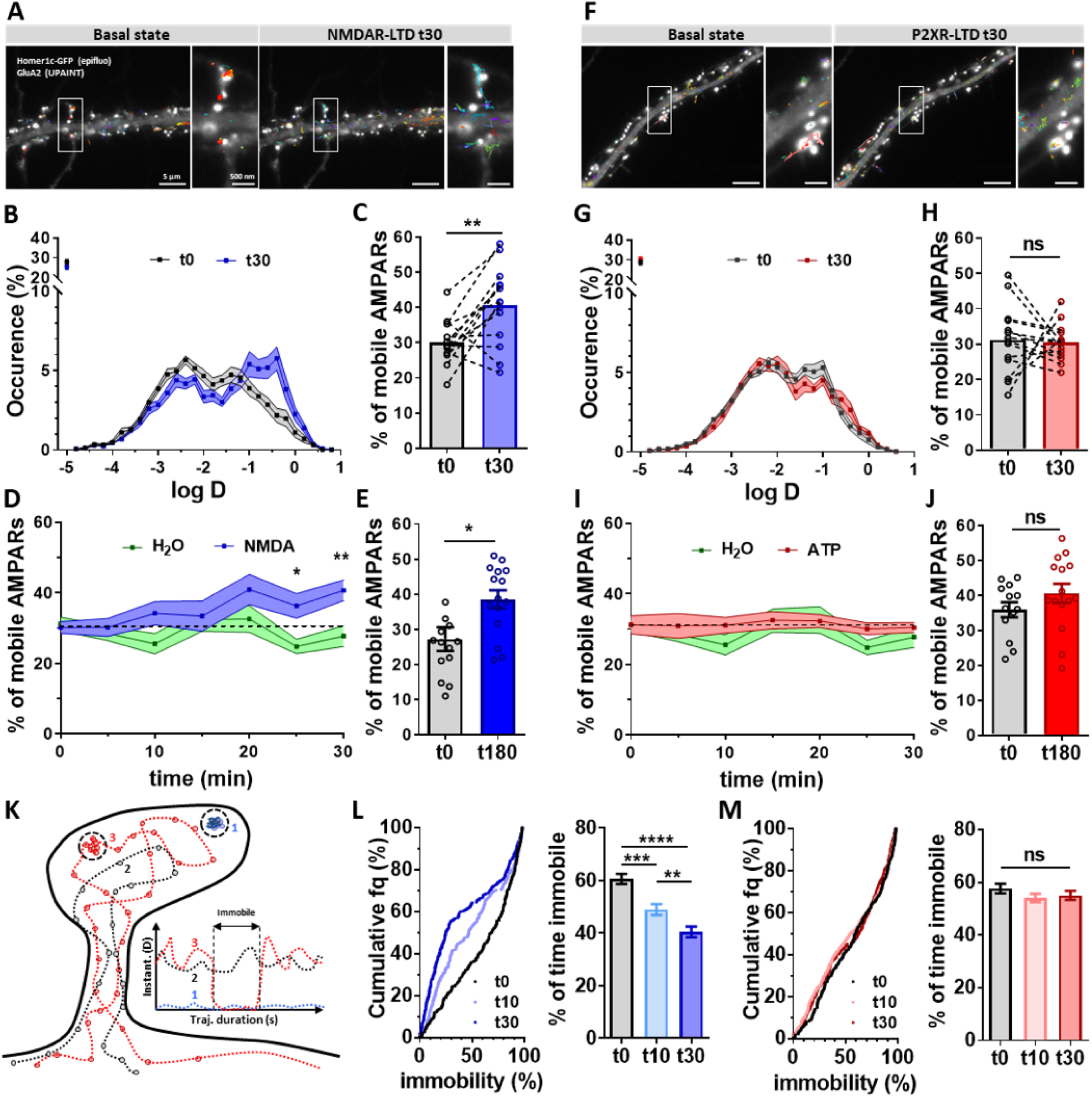
NMDAR-dependent LTD but not P2XR-dependent LTD triggers a long-term increase of AMPAR lateral diffusion. (A) Epifluorescence image of a dendritic segment expressing eGFP-Homer1c as a synaptic marker and GluA2-containing AMPAR trajectories acquired with uPAINT in basal state (left panel) and 30 minutes after NMDA treatment (right panel). (B) Average distribution of the log(D), (D being the diffusion coefficient of endogenous AMPAR) in control condition (black line) and 30 minutes after NMDA treatment (blue line). (C) Average of the mobile fraction per cell, before and 30 minutes after NMDAR-dependent LTD induction (n=14 cells, mean +/- SEM, paired t-test, p=0.0042). (D) Time-lapse (from 0 to 30 minutes) of GluA2-containing AMPAR mobility following NMDAR-dependent LTD induction (blue line) compared to vehicle application (green line) (n=14 and 10 respectively). A significant increase of GluA2-containing AMPAR occurs 25 minutes after NMDA application. (E) Average histograms of the mobile fraction per cell, before and 180 minutes after NMDAR-dependent LTD induction (n=14 and 15 cells, mean +/- SEM, unpaired t-test, p=0.0123). GluA2-containing AMPAR increased mobility remains stable for at least 3 hours. (F to J) Similar experiments as from A to E has been realized with ATP-induced LTD protocol. (F) Epifluorescence image of a dendritic segment expressing eGFP-Homer1c with acquired trajectories of GluA2-containing AMPAR trajectories in basal state (left panel) and 30 minutes after ATP treatment (right panel). (G) Average distribution of the log (D) before (black line) and 30 minutes (red line) after ATP treatment. (H) Average of the mobile fraction per cell extracted from (G), (n=14 cells, mean +/- SEM, paired t-test, p=0.8234). Contrary to NMDA-induced LTD, ATP-induced LTD is not associated with an increase of AMPAR mobility. (I) Time-lapse (from 0 to 30 minutes) of AMPAR mobility following P2XR-dependent LTD induction (red line) compared to vehicle application (green line) (n=14 and 10 respectively). (J) Average histograms of the mobile fraction per cell, before and 180 minutes after P2XR-dependent LTD induction (n=13 and 15 cells, mean +/- SEM, unpaired t-test, p=0.1950). No modification of AMPAR mobility is observed all along the 3 hours experiments. (K) Scheme of the various AMPAR trajectory behaviors. AMPAR can be fully immobile (1, blue line), fully mobile (2 dark line) or alternate between mobile and immobile (3, red line). Calculation of the percentage of immobility all along the trajectory duration give an indication of the avidity of AMPAR for their molecular traps. (L) Variation of the percentage of AMPAR mobility per synaptic trajectories after NMDAR treatment (control (black line), 10 minutes (light blue line) and 30 minutes (dark blue line)). The left panel represents the cumulative distribution and the right panel the mean +/- SEM. (n=252, 235 and 280 synaptic trajectories respectively, one-way ANOVA p<0.0001 and Tukey’s post-test found significant differences p=0.0002 and p<0.0001 between t0 and t10, and t0 and t30 respectively, and p=0.0081 between t10 and t30). (M) Variation of the percentage of AMPAR mobility per synaptic trajectories during ATP-induced LTD (control (black line), 10 minutes (light red line) and 30 minutes (dark red line) following LTD induction). The left panel represents the cumulative distribution and the right panel the mean +/- SEM (n=264, 434 and 326 synaptic trajectories respectively, one-way ANOVA p=0.3360).

As previously described, AMPARs alternate between two main diffusion modes at the plasma membrane: an immobile one when trapped by interaction with scaffolding proteins, and a freely diffusive one (28). We calculated the instantaneous AMPAR diffusion coefficients over time from individual synaptic trajectories (18). For each trajectory, we classified AMPAR movement in three categories: receptors always mobile (class I), receptors always immobile (class II) and receptors alternating between mobile and immobile states (class III) (Figure 2K). Two parameters were computed: the percentage of class II receptors, and the duration of immobilization of the class III receptors. The percentage of fully immobile trajectories was not affected either by NMDA or by ATP treatments at 10 and 30 minutes (data not shown). However, the immobilization duration of mobile receptors, which reflects the avidity of AMPAR for trapping slots, was significantly decreased when LTD was induced by NMDA treatment. Before NMDA application, at synapses, GluA2-containing receptors were immobile 60 percent of the trajectory duration (60.56 +/- 1.91) whereas this duration decreased to 48.92 percent +/- 2.11 after 10 minutes and to 40.37 percent +/-2.03 30 minutes following NMDA treatment (Figure 2L). In contrast, after ATP treatment, this percentage remained unchanged (t0: 57.7 +/- 1.83 percent, t10: 54.14 +/- 1.54 percent, t30: 55.13 +/- 1.71 percent) (Figure 2M). These overall results demonstrated that specifically NMDAR-dependent LTD triggers an increase in AMPAR lateral mobility taking place progressively after the LTD induction phase.

### NMDAR-dependent LTD triggers a depletion of PSD-95 at synapses

Various molecular modifications have been described to impact AMPAR mobility, including a diminution in the number of trapping slots, or a decrease in AMPAR affinity for these slots by modifications in the AMPAR complex composition or phosphorylation status (29). PSD-95 is the main scaffolding protein of the excitatory post-synaptic density and a major actor in AMPAR stabilization at synapses (6,30). Hence, we used dSTORM to measure PSD-95 nanoscale organization following both NMDAR- and P2XR-dependent LTD induction. As previously described, PSD-95 presents two levels of enrichment at synapses (11,12,31); the first one delineating the PSD, the second one corresponding to domains of local concentration into the PSD, beneath the AMPAR nanodomains and facing the glutamate release sites (9–11). Using tesselation-based clustering analysis (32), we extracted the 1st level (termed PSD-95 clusters), and the second level of clustering (termed PSD-95 nanoclusters) (Figure 3A). After NMDA treatment, both PSD-95 clusters and nanoclusters displayed a decrease in number of PSD-95 (estimated number of PSD-95 per clusters t0: 114.7+/-11.6, t10: 82.54+/-10.99, t30: 66.93+/-6.88; per nanoclusters t0: 28.12+/-2.79, t10: 19.91+/-2.02, t30: 18.99+/-2.36) (Figure 3B and C). We also observed a slight decrease in the nanocluster diameter (Figure 3S1). In contrast, following ATP treatment, the number of PSD-95 per clusters or nanoclusters remained unchanged (number of object per PSD-95 clusters t0: 112.5+/-11.25, t10: 104.2+/-9.85, t30: 102.4+/-9.47; per PSD-95 nanoclusters t0: 23.7+/-2.11, t10: 25.13+/-3.25, t30: 21.42+/-1.79) (Figure 3D and E) as their overall organization (Figure 3S1). Altogether, these results indicate that NMDAR-dependent LTD is associated with a change in PSD-95 number and cluster size, which is not the case for P2XR-dependent LTD. This could underlie the observed long-lasting increases in AMPAR mobility in the late phase of NMDAR-dependent LTD.

**Fig. 3.**
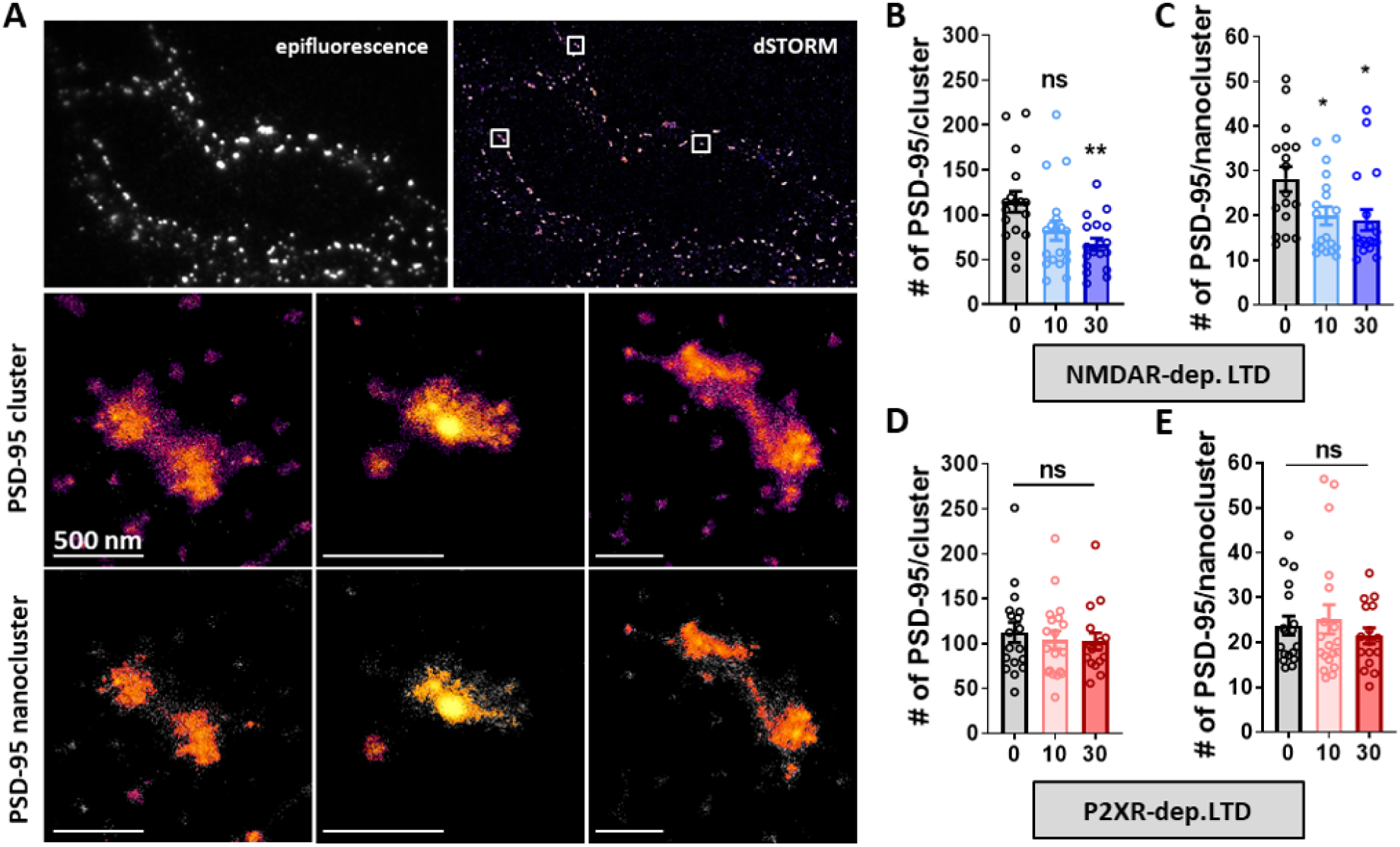
PSD-95 nanocluster organization is modified during NMDA- but not ATP-dependent LTD. (A) Example of endogenous PSD-95 organization along a dendritic shaft (left panel) and in dendritic spines (up-right panel) obtained with dSTORM, represented with SR-Tesseler software. Middle-right panel shows a PSD-95 cluster at the same synapse with an enrichment factor above the average density factor. Down-right panel shows a PSD-95 nanocluster within PSD-95 cluster in the middle-right panel, which corresponds to a PSD-95 structure with a higher density factor than the average PSD-95 cluster’s density. (B and C) Average number of PSD-95 molecules per cluster (B) and per nanoclusters (C) in basal state, 10 and 30 minutes after NMDA treatment (mean +/- SEM, n=17, 19 and 18 respectively, one-way ANOVA p=0.0059 and Dunnett’s post-test found significant between t0 and t30 conditions, p=0.0033 but not between t0 and t10 contions, p=0.0517 for clusters; one-way ANOVA p=0.0189 and Dunnett’s post-test found significant between t0 and t10 and between t0 and t30 conditions, p=0.0348 and p=0.0194 respectively, for nanoclusters). (D and E) Average number of PSD-95 molecules per cluster (B) and per nanoclusters (C) in basal state, 10 and 30 minutes after ATP treatment (mean +/- SEM, n=18, 19 and 16 respectively, one-way ANOVA p=0.7616 and p=0.5269 for clusters and nanoclusters respectively).

### Short-term plasticity is increased during NMDAR-dependent LTD and requires AMPAR lateral diffusion

We have previously established that the pool of mobile AMPARs favors synaptic transmission during high frequency stimulation by allowing desensitized receptors to be replaced by naïve ones (14,18,20). In contrast, a decrease in the proportion of diffusive AMPAR, as triggered by artificial cross-linking (14), CaMKII activation (19) or by associating AM-PARs with the auxiliary proteins TARPs (18) or Shisa6 (21), lead to a significant depression of synaptic transmission during rapid trains of stimulation. Therefore, we investigated whether the increase in AMPAR mobility observed during NMDAR-dependent LTD could affect synaptic responses to high-frequency stimulus trains (5 pulses at 20 Hz). We performed whole-cell patch-clamp recordings of CA1 neurons in acute hippocampal brain slices and measured short-term synaptic plasticity upon stimulation of Schaffer collaterals. LTD was induced by either ATP or NMDA treatment and paired-pulse responses were measured 30 minutes after LTD induction.

We first verified whether both ATP and NMDA treatment triggered a significant decrease in EPSCs amplitude. After NMDA application, we observed a 35 percent decrease in evoked EPSC amplitude (basal state: -37.63+/-2.48 pA, 30’ NMDA: -24.71+/-3.64 pA) and ATP triggered a 28 percent decrease in evoked EPSC amplitude (basal state: -40.89+/- 7.60 pA, 30’ ATP: -29.29+/-7.16 pA) (Figure 4A, B and 4D, E) We then analyzed paired-pulse responses. A representative trace and an average response are shown for NMDA (Figure 4A) and for ATP treatment (Figure 4D). Neurons expressing a P2XR-dependent LTD, which does not trigger an increase in AMPAR mobility, presented a paired-pulse response similar to the one measured during the basal state (Figure 4F). In contrast, neurons treated with NMDA, and thus exhibiting an increase in the proportion of mobile AM-PARs, displayed a significant increase in the paired-pulse ratio compared to those at the basal state (Figure 4C). This increase in the paired pulse ratio was abolished when AMPAR were immobilized by antibody crosslinking (Figure 4S1), as previously described (14,18).

**Fig. 4.**
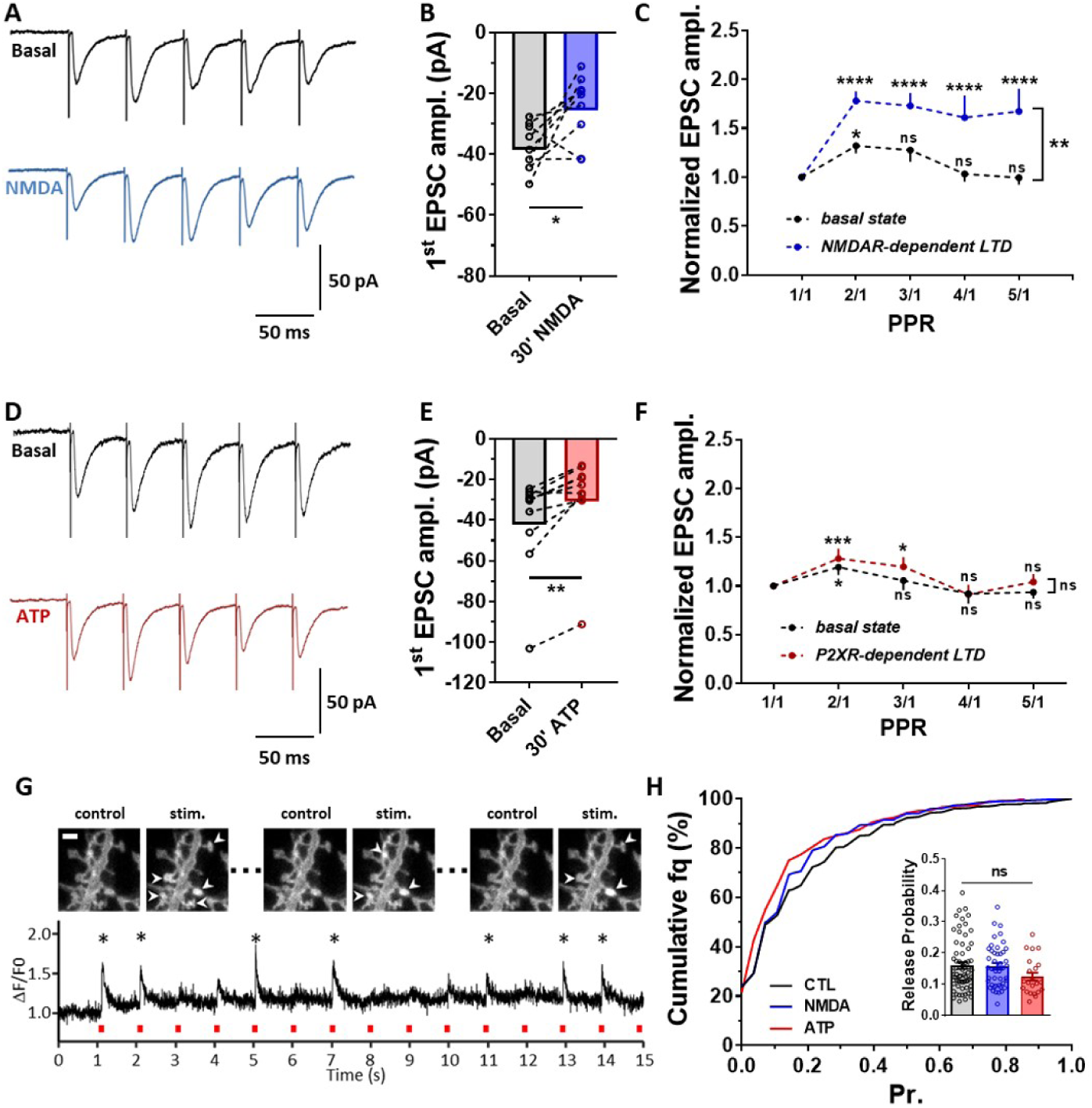
NMDA-dependent LTD is associated to a frequency stimulation facilitation without affecting release probability. (A) Representative traces of synaptic EPSCs in response to 5 stimulations at 20 Hz before (dark line) and 30 minutes after (blue line) NMDAR treatment. (B) Paired-average amplitude of the first response before and 30 minutes after treatment (n=9 cells, mean +/- SEM, paired t-test, p=0.0263). The decrease of the first response demonstrates the efficiency of the LTD protocol. (C) Average of the 5 EPSC amplitudes, normalized by the first response intensity (n=9 cells, mean +/- SEM, two-way ANOVA. For PPR variation, F(4,32)=10.36, p<0.0001, Dunnett’s post-test found significant differences increase of PPR between PPR1/1 and PPR2/1, p=0.0234 at basal state, and between PPR1/1 and either PPR2/1, PPR3/1, PPR4/1 or PPR5/1, p<0.0001, 30 minutes after NMDAR-dependent LTD induction. For basal state vs NMDAR-dependent LTD, F(1,8)=12.85, p=0.0071 and Sidak’s post-test found significant difference between the basal state and 30 minutes after NMDAR-dependent LTD induction for PPR2/1, PPR3/1, PPR4/1 and PPR5/1, p=0.0011, p=0.0013, p<0.0001 and p<0.0001 respectively). A clear facilitation of the currents appears after induction of a NMDAR-dependent LTD. (D, E and F) Similar experiments has been realized when LTD is induced by ATP application, with example of traces in (D). The significant decrease of the first response represented in (E) validate the depression of the synaptic response (n=10 cells, mean +/- SEM, paired t-test, p=0.0012). The average of the 5 responses (F) reveals no facilitation compared to control condition after ATP treatment(n=10 cells, mean +/- SEM, two-way ANOVA. For PPR variation, F(4,36)=7.73, p<0.0001, Dunnett’s post-test found significant differences increase of PPR between PPR1/1 and PPR2/1, p=0.0163 at basal state, and between PPR1/1 and PPR2/1 or PPR3/1, p<0.0004 and p=0.0138 for P2XR-dependent LTD. For basal state vs P2XR-dependent LTD, F(1,9)=1.197, p=0.03023). (G) Example of the fluorescence increase at a synapse expressing iGluSnFR construct during a field stimulation. Responding synapses are labelled with an arrow (upper part). At the bottom, example of the F/F signal obtained at a single synapse. Stars indicate when the synapse is considered as stimulated. (H) Cumulative distribution of the release probability per synapse in control condition (black line) or after LTD induction with either NMDA (Blue line) or ATP (red line) treatment. The mean values per recorded dendrites has been represented in the insert with the same color code. None of the conditions affects significantly the release probability (n=64, 44 and 22 respectively, mean +/- SEM, one-way ANOVA, p=0.1520).

Short-term plasticity has traditionally been attributed to changes in pre-synaptic release probability (33), although it can also arise from AMPAR desensitization (14–17) and be regulated by AMPAR mobility (14,18–20). To decipher between a pre- or post-synaptic origin of the NMDA-induced changes in short term plasticity, we directly measured the pre-synaptic probability of glutamate release before and after NMDAR- or P2XR-dependent LTD using the fluorescent glutamate reporter gene iGluSnFR (34). We expressed iGluSnFR in cultured neurons and measured the variation in post-synaptic fluorescence upon triggering pre-synaptic action potentials by electrical field stimulations (Figure 4G). None of the LTD protocols (ATP or NMDA) changed significantly the pre-synaptic release probability (Figure 4H). These experiments indicate that NMDAR-dependent LTD favors the synaptic responsiveness to high-frequency stimulation through an increase in AMPAR mobility rather than a change in release probability.

### Modeling confirms that increasing AMPARs mobility improves synaptic responsiveness

Both ATP- and NMDA-induced LTD resulted in a decrease in the overall AMPAR number at synapses. Moreover, super-resolution experiments (Figure 2 to 4) show that LTD induced by NMDA, but not by ATP, is associated with an increase in AMPAR mobility. These changes in AMPAR diffusion can be related to a decrease in AMPAR complex affinity for their traps and/or a decrease in the number of synaptic traps (as reported by the decrease of total PSD-95 per synapses) (6,19,30,35,36). To theoretically evaluate the impact of AM-PAR endocytosis or untrapping on AMPAR mobility, organization and synaptic responses, we performed Monte-Carlo simulations using the MCell software (Figure 5). The synaptic shape and perisynaptic environment were obtained from 3D electron microscopy images of hippocampal CA1 stratum radiatum area (9,37–39). The simulation was divided in two sequences. The initial part simulates for 50s, at the ms resolution, the dynamic organization of proteins inside the synapse (Figure 5S1). The second part simulates for 250ms, at the µs resolution, the AMPAR currents following 5 synaptic glutamate releases at 20 Hz (Figure 5B to J, see methods). The protein properties, such as number and diffusion coefficient, were implemented into the model based on the results obtained with super-resolution imaging techniques and in agreement with previous papers (40), i.e., 200 PSD-95 and 120 AMPAR molecules (half in an internal pool, half at the surface). Interactions between proteins were implemented following the scheme (Figure 5A), and affinity constants (k) were adjusted to reach, at the equilibrium, a distribution similar to the one observed by microscopy (Figure 5S1). Based on the literature and our experimental results, we tested the relation between simulated AMPAR current amplitudes and variations of three different interaction constants: the endocytosis rate (k8*3), (ii) the affinity of AMPAR for PSD-95 (trapping rate; k4*4) and (iii) the removal of PSD-95 into the PSD named PSD-95 inactivation rate (k6*4). The 3 times increase in the endocytosis rate (noticed k8*3Endo) which could correspond to the initial phase of both ATP- and NMDA-induced LTD, triggered a 25 percent decrease in the number of activated AMPAR (Figure 5B). This value is similar to the current amplitude decrease measured with electrophysiology. In parallel, the increase in AMPAR untrapping (k4*4Untrap) and the increase in PSD-95 inactivation (k6*4Inact) led respectively to a 32 percent and 30 percent decrease in AMPAR current amplitude at the first glutamate release (Figure 5E and 5H). Interestingly, these two modifications triggered a net increase in the number of mobile AMPAR (as illustrated for k6*4 in Figure 6S1B, blue line), mimicking the results observed during the late phase of NMDAR-dependent LTD (30 minutes and 3 hours after LTD induction). In another set of modeling experiments, we induced an AMPAR depletion into domains by increasing endocytosis rate (k8*3) and then put back this rate at its initial value of 1. A rapid replenishment of nanodomains is observed (Figure 5SA). Interestingly, this replenishment could be counterbalanced by an increase of AMPAR untrapping (k4*4) (Figure 5S2B). We then determined the synaptic responses following trains of 5 stimulations at 20 Hz in these various conditions. K4*4Untrap and k6*4Inact conditions triggered similar increase in paired pulse ratio, with a 20.2 and 18.7 percent increase in AMPAR activation respectively for the second release and a 25.7 and 20.4 percent for the third one (Figure 5F,G,I and J). While the k8*3Endo condition, which does not impact AMPAR mobility, did not modify the synaptic response in frequency (Figure 5B). Using a realistic model of AMPAR organization, these simulation confirm that an increase in the pool of freely diffusing AMPAR induced by either a decrease of their affinity for traps or a decrease in the number of traps, is sufficient to trigger a paired-pulse facilitation similar to the one observed on brain slices by electro-physiological experiments.

**Fig. 5.**
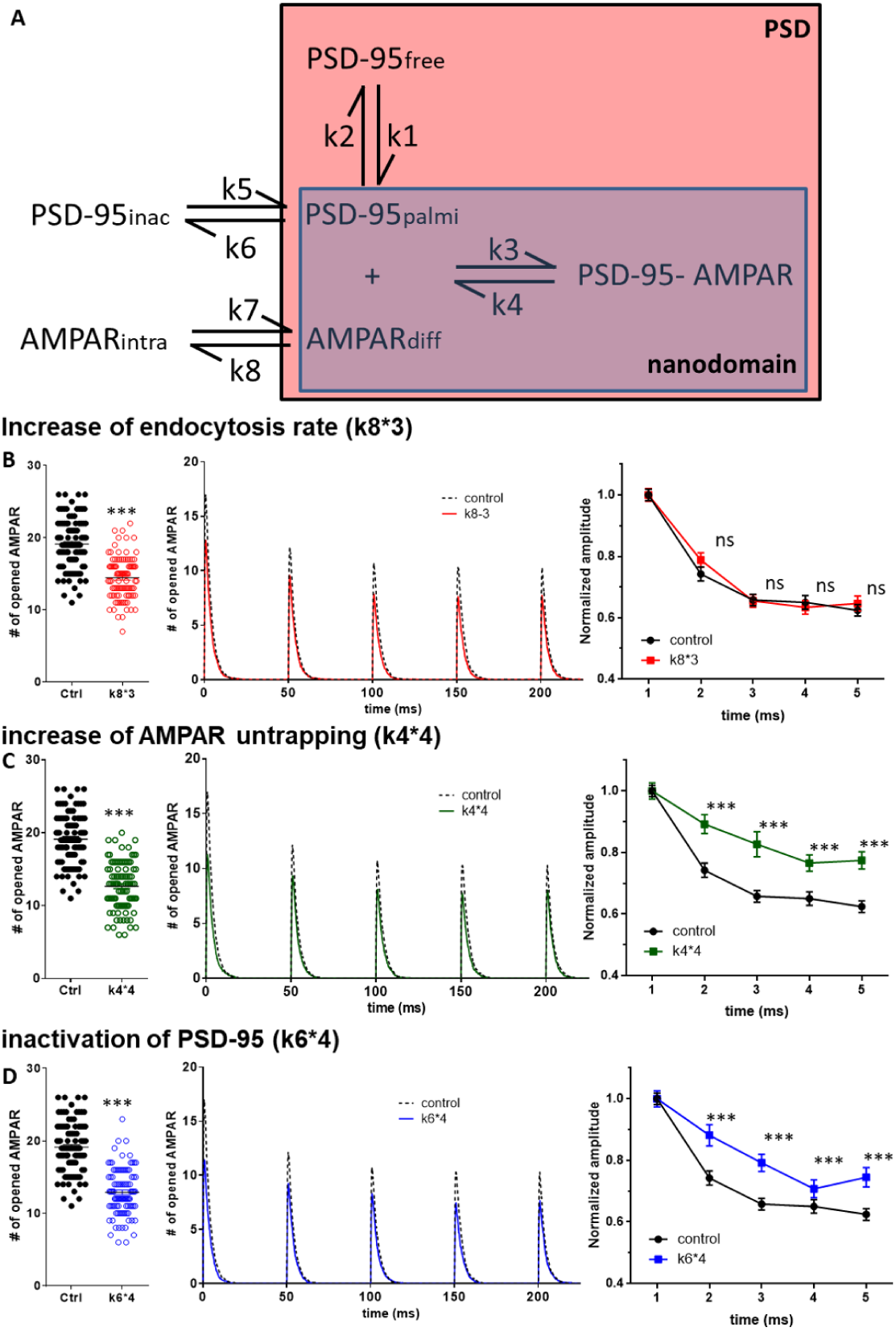
In silico simulations confirm that AMPAR untrapping induced both a depression of synaptic currents and an increase of synaptic responsiveness. (A) Representation of the main interactions which define the AMPAR organization/ trapping. PSD-95 can diffuse freely, be slowly mobile and confined into nanodomain when palmitoylated, or be inactivated. AMPAR can be endocytosed, freely mobile at the surface or being trapped by palmitoylated PSD-95 into the domain. Three various kinetic rate constant modifications trigger a synaptic depression, (1) an increase of endocytosis rate, mimicking the initial phase of the NMDAR-dependent LTD and the P2XR-dependent LTD (B). (2) an increase dissociation between palmitoylated PSD-95 and AMPAR (C), triggering an increase of the pool of diffusive AMPAR. (3) A decrease of the total number of PSD through an increase of their inactivation rate(D), mimicking the depletion of PSD-95 observed during NMDAR-dependent LTD. For each condition, we report in the right panel, the number of open AMPARs during the first glutamate release, before (dark dots) and after (color dots) modification of the parameter. A significant decrease of AMPAR response similar to the depression experimentally measured is observed in all conditions. Middle panel, the average traces of the equivalent AMPAR current following 5 glutamate releases at 20 Hz. Right panel, the average of the AMPAR equivalent current following the 5 releases, normalized by the initial response. 100 independent simulations are realized in each condition. When depletion of synaptic AMPAR are induced by increasing the endocytosis, there is no modification of simulated paired pulse ratio while AMPAR untrapping and PSD-95 inactivation conditions triggers both a significant increase of PPR.

Altogether, these experiments indicate that long-term synaptic depression-induction led to a depletion of AMPAR into synaptic nanodomains. This reorganization can be mainly attributed to the depletion of the pool of surface receptor most probably through an increase of endocytosis rate and occurs following various LTD induction paradigms. However, our experiments reveal that only following the induction of the NMDAR-dependent LTD, a more complex synaptic reorganization occurs, dictated by a progressive reorganization of PSD-95 and triggering an increase of AMPAR mobility. This change of the proportion of diffusive receptor is correlated with an increase of synaptic responsiveness following high frequency stimulation. Experimental observations are confirmed by modeling, which show that an increase in the mobile fraction of AMPAR is sufficient to trigger a paired-pulse facilitation. We demonstrated here that LTD cannot be restricted to a simple decrease in synaptic AMPAR by endocytosis. Independent LTD protocols differently affect AMPAR nanoscale organization and dynamic, and each type of LTD present various temporal phases rely to specific molecular reshuffling.

## Discussion

Using single molecule localization-based super-resolution microscopy on live and fixed neurons, combined with electrophysiology and modeling, we characterized the nanoscale modifications in AMPAR organization and dynamic triggered by two different types of LTD-inducing stimuli and estimated their impact on frequency dependent synaptic current properties. We identified a common induction phase going through a depletion of AMPAR content both in nanodomains and at synapses, leading to a decrease in synaptic strength. In a subsequent phase, NMDAR-dependent LTD is associated with a net increase in the proportion of mobile AMPAR, and a depletion in PSD-95 clusters. Importantly, our experimental data and simulations using a realistic model indicate that this change in AMPAR dynamic allows synapses to improve their responsiveness to high frequency stimulations. Altogether, our data introduce a new level of synaptic integration, where various LTD types do not similarly impact on synaptic responses. This argues for a new mechanism through which regulation of AMPAR surface density and diffusion following specific post-synaptic signaling to trigger LTD allows to adjust the capacity of synapses to encode pre-synaptic activity.

### P2XR- and NMDAR-dependent LTD are associated with nanodomain depletion proportional to the decrease in AMPAR mEPSC amplitudes

Since the discovery that AMPARs are organized in nanodomains (11,12,31) and that AMPAR clusters are aligned with pre-synaptic release sites (9,10), our view of synapse function, and how it could be plastic, has much evolved. Together with previous findings (13,14,41,42), this has introduced the concept that post-synaptic plasticity could arise not only from absolute changes in AMPAR content, but also from their local reorganization regarding glutamate release site at the nanoscale. Thus, it became important to revisit LTD through this new prism. AMPAR-mediated current amplitude changes observed during LTD could arise from (i) a modification of the domain structure leading to a decrease in the packing of receptor (13); (ii) a misalignment between the pre- and the post-synaptic machinery (9,42) or more simply (iii) a depletion of the domains and overall decrease in synaptic AMPAR content. Here we report that both NMDA and ATP induce a synaptic depression based, at least partially, on post-synaptic nanoscale modifications. Super-resolution experiments revealed no modification of the overall nanodomain dimensions, and no pre-post synaptic misalignment. However, we observed a rapid (less than 10 minutes) depletion in the number of AMPAR per nanodomain. Interestingly, the extent of synaptic AMPAR depletion (around 25 percent) is proportional to the extent of mEPSC depression. This suggests that synaptic depression can be mainly attributed to a decrease in post-synaptic AMPAR content.

### The late phases of P2XR- and NMDAR-dependent LTD are distinct

Both P2XR- and NMDAR-dependent LTD induction have been attributed to a net increase in AMPAR endocytosis rate (24,43–45). However, the mechanisms involved in the longterm maintenance of synaptic depression are less clear, and as demonstrated with simulation (Figure 5S2), a transient increase in endocytosis rate triggers only a transient depression and not a long term one. In principle, 3 variables can be listed to explain the maintenance in the depression of AMPAR number at synapses following LTD induction: the maintenance of an endocytosis rate higher than exocytosis; a decrease in the number of trapping slots at the PSD, or a decrease in the affinity of AMPAR complex for these trapping slots. In previous studies, we demonstrated that ATP treatment triggers a large increase in endocytosis both in Xenopus oocytes and in neurons (24). This is due to activation of CAMKII, which phosphorylates AMPAR at the Ser567 and Ser831 on GluA1 and GluA2 subunits (26). Our modeling studies indicate that an increase in endocytosis is sufficient to decrease the AMPAR synaptic content (Figure 5), but that a return of the endocytosis rate to its initial level leads to a rapid replenishment of AMPAR at the synapse (Figure 5S2A). Here, we demonstrate (figure 1) that P2XR-dependent LTD is stable for at least 3 hours. As we could not detect an ATP induced modification in PSD-95 organization or in AMPAR affinity for trapping slots, we hypothesize that the initial unbalance in endocytosis/exocytosis ratio responsible of LTD induction is maintained to support the long-term maintenance of LTD. However, we cannot determine whether a long term endocytosis/exocytosis unbalance is due to the maintenance of a high endocytosis rate or to a decrease of exocytosis rate as already described after NMDA treatment by Fujii et al. (46). In parallel, a large increase in the AMPAR endocytosis rate after NMDA treatment has been reproducibly reported (43–45). This increase in AMPAR internalization rate, likely responsible for the initiation of the LTD, stays for less than 10 minutes (45). Contrary to ATP treatment, the maintenance of the depression is associated with changes of both AMPAR mobility and a decrease of the number of PSD-95 slots. These modifications likely happen via series of phosphorylations/ dephosphorylations of the AMPAR or its auxiliary subunits (47,48). Simulations (Figure 5S2) confirmed that a decrease in AMPAR affinity for synaptic traps succeed to maintain the depression by compensating for the decrease of endocytosis rate after ten minutes. Our work suggests that NMDA- and ATP-induced LTD are mediated by distinct molecular mechanisms that both maintain AMPAR depression. After a similar induction phase likely due to an increase in AMPAR endocytosis rate, we hypothesized that the activation of P2XR should maintain an endocytosis/exocytosis unbalance, whereas NMDAR activation triggers a more profound re-organization of the synapse via a decrease in AMPAR affinity for the synaptic traps, and/ or a decrease of PSD-95 in the domains.

### A decrease in AMPAR trapping at domains takes place gradually when NMDAR-dependent LTD is induced

The existence of various phases during NMDAR-dependent LTD has been described previously. Rosendale et al. demonstrated that the endocytosis rate increases immediately after LTD induction but then goes back to its original value after approximately 10 minutes (45). Sanderson et al. 2016, reported a rapid and transient increase in calcium-permeable AMPARs (49). Here, we observed a delayed increase in the proportion of mobile AMPAR, developing in the half hour after LTD induction, and a depletion of PSD-95 both inside the nanocluster and inside the entire PSD. Two main mechanisms could explain this change in AMPAR dynamic organization. During synaptic long-term potentiation (LTP), the phosphorylation state of TARPs is modified, directly affecting the affinity of AMPAR complex for the synaptic traps (19,30,47). Thus, during NMDAR-dependent LTD, activation of calcineurin and other phosphatases leading to AMPAR and TARPs dephosphorylation could affect their stabilization at synapses and participate to the observed increase in AMPAR mobility (47,50,51). Alternatively, a decrease in the number of traps corresponding to the decrease in PSD-95 we measured could also lead to an increased AMPAR mobility. We have tested these two hypotheses via modeling, hypothesis 1 being the k4*4 conditions and the hypothesis 2 the k6*4 one (Figure 5), and concluded to a similar increase of AMPAR mobility and synaptic depression.

### The AMPAR mobility increase induced during NMDAR-dependent LTD mediates an improvement in synaptic responsiveness

Short term synaptic plasticity (STP) depends on many factors, including pre-synaptic transmitter release (33), post-synaptic AMPAR desensitization (14–17) and AMPAR surface diffusion (14,18–20). The latter mechanism potentiates synaptic responses to sequential stimuli by allowing desensitized receptors to be exchanged by naïve ones, hence improving the rate of recovery of synaptic depression due to AMPAR desensitization. Strikingly, we observed that following NMDA treatment, but not ATP, the ability of synapses to follow high frequency stimulation is improved. This differential effect of NMDA and ATP on STP mirrors their effect on AMPAR mobility. Therefore, it is attractive to suggest that NMDA treatment increases STP through the increase in AMPAR mobility. This hypothesis is supported by the observation that: (i) NMDA does not modify pre-synaptic release probability, (ii) the time course of STP potentiation parallels that of the NMDA-induced increase in mobility, i.e. both processes only happen 30 minutes after NMDAR-dependent LTD, (iii) blocking the NMDA-induced increase in AMPAR mobility by AMPAR X-linking prevents the potentiation of STP, (iv) P2XR-dependent LTD, which does not affect AMPAR mobility, does not modify STP and (v) modeling confirmed that increasing the AMPAR mobile pool by decreasing their affinity for the traps or the number of traps, favors the ability of synapses to follow high frequency stimulation. Of note, a similar increase in the paired pulse ratio upon an increase in the proportion of mobile AMPAR has already been observed upon digestion of the extracellular matrix (20). Here, for the first time, our data suggest that a physiological increase in AMPAR mobility, after a protocol that induces LTD, triggers an improvement of synaptic responsiveness through potentiation of STP. This induced reshuffling of PSD-95 and AMPAR nanoscale organization induced by NMDAR-dependent LTD clearly demonstrates that synaptic depression does not solely correspond to a decrease in the synaptic response amplitude but to deeper changes at the nanoscale which modify the capacity of the depressed synapses to encode pre-synaptic inputs.

## ETHICAL APPROVAL

All experiments were approved by the Regional Ethical Committee on Animal Experiments of Bordeaux.

## AUTHOR CONTRIBUTIONS

B.C performed all dSTORM and single-particle tracking experiments, culture and slice electrophysiology. M.M performed iGluSnfr experiments and D.P. developed the software to analyze iGluSnfr experiments. E.H, T.M.B and T.J.S created the MCell model and E.H performed the simulations. C.B, A.K and J.B.S developed the super-resolution analysis software. R.V.K and A.B.S participate to conception and validation of some hypothesis and writing of the paper. E.H conceived and supervised the study. D.C financed the study. E.H, B.C and D.C wrote the manuscript. All authors contributed to the preparation of the manuscript.

## ACKNOWLEDGEMENTS

We acknowledge E.Gouaux for the anti-GluA2 antibody and J.S. Marvin for the iGluSnFR plasmid; the Bordeaux Imaging Center, part of the FranceBioImaging national infrastructure (ANR-10INBS-04-0, for support in microscopy; E.Normand and the IINS in vivo facility for animal husbandry. We thank the IINS cell biology core facilities (LABEX BRAIN [ANR-10-LABX-43]) and in particular C. Breillat, E. Verdier, and N. Retailleau for cell culture and plasmid production, and Jorge Aldana from the Salk institute for computing support. This work was supported by funding from the Ministère de l’Enseignement Supérieur et de la Recherche (ANR NanoDom and AMPAR-T), Fulbright and Philippe foundation to E.H. and D.C., Centre National de la Recherche Scientifique (CNRS), ERC grant ADOS (339541) and DynSynMem (787340) to D.C., Fondation pour la Recherche Médicale fellowship to B.C.

## Supplementary information

### Material and Methods

#### Hippocampal neuron culture and transfection

Culture are realize as described in Nair et al. 2013. For uPAINT experiments, neurons were electroporated (4D-Nucleofector system, Lonza, Switzerland) just after dissection with eGFP-Homer1c, otherwise, neurons are transfected at DIV 10 with calcium phosphate

#### Sample preparation and immuno-labeling

For dSTORM and confocal imaging of PSD-95, primary neuronal cultures were treated either with 30 µM NMDA (Tocris) for 3 minutes or with 100 µM ATP (Sigma-aldrich) for 1 minute and fixed with PFA 10 or 30 minutes after. PFA was quenched with NH4Cl 50 mM for 10 minutes. A permeabilization step with 0.2 percent triton X100 for 5 minutes was performed. Cells were washed 3 times for 5 min in 1x PBS. After 3 washes with 1x PBS, unspecific staining was blocked by incubating coverslips in 1 percent BSA for 1h at room temperature. Cells were then incubated with monoclonal mouse anti-PSD-95 antibody (MA1-046, ThermoFischer), diluted in 1 percent BSA at 1/500, at room temperature for 1 hour. Coverslips were rinsed 3 times in 1 percent BSA solution and incubated in 1 percent BSA for 1h at room temperature. Primary antibodies were revealed with Alexa 647 (dSTORM) or Alexa 488 (confocal) coupled anti-mouse IgG secondary antibodies (ThermoFisher, A21235 and A11001).

For dSTORM imaging of pre- to post-synaptic alignment, primary neuronal cultures were incubated 0, 10 or 30 minutes after NMDA treatment with monoclonal mouse anti-GluA2 antibody 11 for 7 minutes at 37° C and then fixed with 4 percent PFA. After permeabilization, unspecific staining was blocked by incubating coverslips in 1 percent BSA for 1h at room temperature. Cells were then incubated with a polyclonal rabbit anti-RIM 1/2 antibody (synaptic systems, 140 203). Primary antibodies were revealed with Alexa 532 coupled anti-mouse IgG secondary antibodies (ThermoFisher, A21235) and with Alexa 647 coupled anti-rabbit IgG secondary antibodies (ThermoFisher, A21244). For dSTORM imaging of GluA2-containing AMPARs, primary neuronal cultures were treated with 30 µM NMDA (Tocris) for 3 minutes 23 or with 100 µM ATP (Sigma-aldrich) for 1 minute in presence of CGS15943 (3 µM) 24,26. After 10 or 30 minutes, neurons were incubated with a monoclonal mouse anti-GluA2 antibody (mouse antibody, provided by E. Gouaux, Portland, USA, 9,11,18 for 7 minutes at 37°C and then fixed with 4 percent PFA. Then, cells were washed 3 times for 5 min in 1x PBS. PFA was quenched with NH4Cl 50 mM for 10 minutes. Unspecific staining was blocked by incubating coverslips in 1 percent BSA for 1h at room temperature. Primary antibodies were revealed with Alexa 647 coupled anti-mouse IgG secondary antibodies (ThermoFisher, A21235).

For dSTORM and confocal imaging of PSD-95, primary neuronal cultures were treated either with 30 µM NMDA (Tocris) for 3 minutes or with 100 µM ATP (Sigma-aldrich) for 1 minute and fixed with PFA 10 or 30 minutes after. PFA was quenched with NH4Cl 50 mM for 10 minutes. A permeabilization step with 0.2 percent triton X100 for 5 minutes was performed. Cells were washed 3 times for 5 min in 1x PBS. After 3 washes with 1x PBS, unspecific staining was blocked by incubating coverslips in 1 percent BSA for 1h at room temperature. Cells were then incubated with monoclonal mouse anti-PSD-95 antibody (MA1-046, ThermoFischer), diluted in 1 percent BSA at 1/500, at room temperature for 1 hour. Coverslips were rinsed 3 times in 1 percent BSA solution and incubated in 1 percent BSA for 1h at room temperature. Primary antibodies were revealed with Alexa 647 (dSTORM) or Alexa 488 (confocal) coupled anti-mouse IgG secondary antibodies (ThermoFisher, A21235 and A11001).

For dSTORM imaging of pre- to post-synaptic alignment, primary neuronal cultures were incubated 0, 10 or 30 minutes after NMDA treatment with monoclonal mouse anti-GluA2 antibody 11 for 7 minutes at 37° C and then fixed with 4 percent PFA. After permeabilization, unspecific staining was blocked by incubating coverslips in 1 percent BSA for 1h at room temperature. Cells were then incubated with a polyclonal rabbit anti-RIM 1/2 antibody (synaptic systems, 140 203). Primary antibodies were revealed with Alexa 532 coupled anti-mouse IgG secondary antibodies (ThermoFisher, A21235) and with Alexa 647 coupled anti-rabbit IgG secondary antibodies (ThermoFisher, A21244).

#### direct STochastic Optical Reconstruction Microscopy (dSTORM)

dSTORM experiments were done on fixed immunolabeled neurons. dSTORM imaging was performed on a LEICA DMi8 mounted on an anti-vibrational table (TMC, USA), using a Leica HCX PL APO 160x 1.43 NA oil immersion TIRF objective and fibber-coupled laser launch (405 nm, 488 nm, 532 nm, 561 nm and 642 nm) (Roper Scientific, Evry, France). Fluorescent signal was collected with a sensitive EMCCD camera (Evolve, Photometrics, Tucson, USA). The 18 mm coverslips containing neurons were mounted on a Ludin chamber (Life Imaging Services, Switzerland) and 600 µL of imaging buffer was added 52. Another 18 mm coverslip was added on top of the chamber to minimize oxygen exchanges during the acquisition to limit contact with the oxygen of the atmosphere. Image acquisition and control of microscope were driven by Metamorph software (Molecular devices, USA). Image stack contained typically 40,000-80,000 frames. Selected ROI (region of interest) had dimension of 512×512 pixels (one pixel = 100 nm). The power of the 405 nm laser was adjusted to control the density of single molecules per frame, keeping the 561 laser intensity constant. Multi-color fluorescent microspheres (Tetraspeck, Invitrogen) were used as fiducial markers to register long-term acquisitions and correct for lateral drifts. Super-resolution images with a pixel size of 25 nm were reconstructed using WaveTracer software 53 operating as a plugin of MetaMorph software.

#### Cluster analysis

##### AMPAR nanodomain analysis

Localization of Alexa-647 signals was performed using PalmTracer, a software developed as a MetaMorph plugin by J.B. Sibarita group (Interdisciplinary Institute for Neuroscience). AMPAR nanodomain properties were extracted from super-resolution images corrected for lateral drift as described in previous studies 11,18.

##### PSD-95 cluster analyses

PSD-95 clusters and nanoclusters were then identified using SR-Tesseler software 32. A first automatic threshold of normalized density DF = 1 was used to extract clusters of PSD-95 (cluster of level 1) having an enrichment factor higher than the average localization density, corresponding to Post-Synaptic Densities (PSD). A second threshold of DF = 1 applied on the localizations inside these clusters was used to identify the PSD-95 nanoclusters corresponding to domains.

##### AMPAR-RIM1/2 cluster distance measurement

Localizations of Alexa-532 and Alexa-647 were corrected for chromatic aberration using a correction matrix calibrated from a set of tetraspeck beads imaged both with 642nm and 532nm excitation wavelengths. Clusters of AMPARs and clusters of RIM1/2 proteins were detected using the multicolor version of SR-Tesseler software as described previously for single color SR-Tesseler, and distances between clusters detected in each color were measured within each synaptic ROI in order to solely measure the distances between objects belonging to the same synaptic contact.

#### universal Point Accumulation Imaging in Nanoscale Topography (uPAINT)

For LTD experiments lasting 30 minutes, chemical treatments to induce LTD were added into the Ludin chamber after the first movie acquisition. NMDAR-dependent LTD was induced using NMDA (Tocris Bioscience) at 30µM for 3 minutes, while P2XR-dependent LTD was induced using ATP (Sigma-Aldrich) at 100µM for 1min in presence of CGS15943 (3µM) as described in Pougnet et al. 2014 24. Imaging solution was washed and replace by new Tyrode solution and ATTO647N coupled-anti-GluA2 antibody at low concentration was added. A 4000 frame movie of the same dendritic segment was recorded at 50Hz every 5 minutes for 30 minutes.

LTD experiments lasting 3 hours were performed in non-paired conditions. Some coverslips were treated with the chemical compound inducing LTD while control are treated with water. Coverslips were placed into the culture dish at 37°C and 5percent CO2 and were imaged as previously described for uPAINT experiments. A single movie of 4000 frames at 50Hz was acquired. For u-PAINT experiments, the 18 mm coverslip containing neurons was mounted on a Ludin chamber (Life Imaging Services, Switzerland). Cells were maintained in a Tyrode solution equilibrated at 37°C and composed of the following (in mM): 15 D-Glucose, 100 NaCl, 5 KCl, 2 MgCl2, 2 CaCl2, 10 HEPES (pH7.4; 247mOsm). Imaging was performed on a Nikon Ti-Eclipse microscope equipped with an APO 100x 1.49 NA oil immersion TIRF objective and laser diodes with following wavelength: 405 nm, 488 nm, 561 nm and 642 nm (Roper Scientific, Evry, France). A TIRF device (Ilas, Roper Scientific, Evry, France) was placed on the laser path to modify the angle of illumination. Fluorescence signal was detected with sensitive EMCCD camera (Evolve, Roper Scientific, Evry, France). Image acquisition and control of microscope were driven by Metamorph software (Molecular devices, USA). The microscope was caged and heated in order to maintain the biological sample at 37°C. The first step consisted to find an eGFP-Homer1c transfected neuron. This construct was used in order to visualize the neuron of interest and the synaptic area for more synaptic trajectory analysis. After selection of the dendritic segment of interest, ATTO647N coupled-anti-GluA2 antibody (mouse antibody, provided by E. Gouaux, Portland, USA) at low concentration was added in the Ludin chamber to sparsely and stochastically label endogenous GluA2-containing AMPARs at the cell surface. The TIRF angle was adjusted in oblique configuration to detect ATTO647N signal at the cell surface and to decrease background noise due to freely moving ATTO647N coupled antibodies. 647nm laser was activated at a low power to avoid photo-toxicity but allowing a pointing accuracy of around 50nm, and 4000 frames at 50Hz were acquired to record AMPAR lateral diffusion at basal state.

For LTD experiments lasting 30 minutes, chemical treatments to induce LTD were added into the Ludin chamber after the first movie acquisition. NMDAR-dependent LTD was induced using NMDA (Tocris Bioscience) at 30µM for 3 minutes, while P2XR-dependent LTD was induced using ATP (Sigma-Aldrich) at 100µM for 1min in presence of CGS15943 (3µM) as described in Pougnet et al. 2014 24. Imaging solution was washed and replace by new Tyrode solution and ATTO647N coupled-anti-GluA2 antibody at low concentration was added. A 4000 frame movie of the same dendritic segment was recorded at 50Hz every 5 minutes for 30 minutes.

LTD experiments lasting 3 hours were performed in non-paired conditions. Some coverslips were treated with the chemical compound inducing LTD while control are treated with water. Coverslips were placed into the culture dish at 37°C and 5percent CO2 and were imaged as previously described for uPAINT experiments. A single movie of 4000 frames at 50Hz was acquired.

#### Single-Particle Tracking analysis

Single molecule localization, tracking and Mean Square Displacement (MSD) of ATTO-647N signals (uPAINT) were computed using PALMTracer software like in Nair et al 2013. From the MSD, two parameters were extracted: (i) the diffusion coefficient (D) corresponding to the global diffusion of the trajectory were calculated by linear fit of the first four points of the Mean Square Displacement plots. (ii) The instantaneous diffusion, corresponds to the variations of the D values all along the trajectory duration (see 18).

#### Electrophysiological recordings

##### mEPSC recordings in neuronal culture

Coverslips of eGFP-Homer1c electroporated neurons were placed in a Ludin Chamber on an inverted motorized microscope (Nikon Eclipse Ti) and transfected neurons were identified under epifluorescence from the GFP signal. Extracellular recording solution was composed of the following (in mM): 110 NaCl, 5.4 KCl, 1.8 CaCl2, 0.8 MgCl2, 10 HEPES, 10 D-Glucose, 0.001 Tetrodotoxin and 0.05 Picrotoxin (pH 7.4; 245 mOsm/L). Patch pipettes were pulled using a horizontal puller (P-97, Sutter Instrument) from borosilicate capillaries (GB150F-8P, Science Products GmbH), and parameters are adjusted to reach a resistance of 4-6 M?. The pipettes are filled with intracellular solution composed of the following (in mM): 100 K-gluconate, 10 HEPES, 1.1 EGTA, 3 ATP, 0.3 GTP, 0.1 CaCl2, 5 MgCl2 (pH 7.2; 230 mOsm). Recordings were performed using an EPC10 patch clamp amplifier operated with Patchmaster software (HEKA Elektronik). Whole-cell voltage clamp recordings were performed at room temperature and at a holding potential of -70mV. Unless specified otherwise, all chemicals were purchased from Sigma-Aldrich except for drugs, which were from Tocris Bioscience. Miniature EPSC analysis were performed using a software developed by Andrew Penn, the matlab script is available on MATLAB File Exchange, ID: 61567; http://uk.mathworks.com/matlabcentral/fileexchange/61567-peaker-analysis-toolbox.

##### Paired-Pulse Response recordings in acute slices

Acute slices were prepared from P16-18 Sprague-Dawley rats of both sexes. Rats were anesthetized with 5percent isofluorane prior to decapitation according to the European Directive rules (2010/63/EU). Brain were quickly extracted and the two hemispheres were separated and placed in ice-cold, oxygenated (95percent O2,5 percent CO2) sucrose-based artificial cerebrospinal fluid (ACSF) containing (in mM): 250 Sucrose, 2 KCl, 7 MgCl2, 0.5 CaCl2, 11 Glucose, 1.15 NaH2PO4 and 26 NaHCO3 (pH 7.4; 305 mOsm/L). Sagittal slices were cut (350 µm thick) and incubated for 30 minutes at 32°C in carbogenated ACSF (95percent O2,5percent CO2) containing (in mM): 126 NaCl, 3.5 KCl, 2 CaCl2, 1 MgCl2, 1.2 NaH2PO4, 25 NaHCO3 and 12.1 Glucose (pH 7.4; 310 mOsm/L). Subsequently, slices were incubated for 30 minutes at room temperature and used until 5 hours after preparation. Experiments were performed in a submerged recording chamber at 30-32°C with continuous perfusion of carbogenated ACSF added with Gabazine (2µM) and CGP52432 (2µM). The intracellular solution was composed of (in mM): 130 Cs methane sulfonate, 10 HEPES, 10 EGTA, 2 MgCl2, 1 CaCl2, 4 Na2-ATP, 0.4 Na-GTP and 5 QX314. Synaptic responses were obtained by 5 stimulations of Schaffer collateral with 0.2 ms pulses at 50 Hz. 20 series spaced by 20 seconds were performed. LTD was induced by perfusion of NMDA (30µM, 3min) or ATP (100µM, 1min), in presence of CGS15943 3µM. Another 20 series of 5 stimulations at 50HZ were performed 30min after LTD induction. Average of each 20 series were calculated. Each response was normalized to the first one. Paired-Pulse Ratios were measured using Stimfit software taking into account fully successful paired-pulse response (trials with failures were rejected from analysis).

#### iGluSnFR imaging and release probability measurement

Transfection of iGluSnFR (Marvin et al., 2013) was performed on banker neuronal culture at 6 days in vitro (DIV) by a calcium phosphate transfection procedure. Experiments were carried out at 15-18 DIV. Thirty minutes after induction of LTD or water application for control, the neuronal preparation was placed under continuous perfusion in a Tyrode solution containing 100mM NaCl, 5mM KCl, 2mM CaCl2, 2mM MgCl2, 15mM glucose, 10mM HEPES pH 7.4. Experiments were performed at 35°C on an inverted microscope (IX83, Olympus) equipped with an Apochromat N oil ×100 objective (NA 1.49). Samples were illuminated by a 473-nm laser (Cobolt) and emitted fluorescence was detected after passing a 525/50nm filter (Chroma Technology Corp.). Images were acquired at a resolution of 100 × 100 pixels every 3 ms with a sCMOS camera (Prime 95B; Photometrics) controlled by MetaVue7.1 (Roper Scientific). Neurons were stimulated by electric field stimulation (platinum electrodes, 10mm spacing, 1ms pulses of 50mA and alternating polarity at 1 or 10Hz) applied by constant current stimulus isolator (SIU-102, Warner Instruments) in the presence of 10µM 6-cyano-7-nitroquinoxaline-2,3-dione (CNQX) and 50µM d,l-2-amino-5-phosphonovaleric acid (AP5) to prevent recurrent activity. Image analysis was performed with custom-written macros in MATLAB (MathWorks) using an automated detection algorithm. At the end of the experiments, a 10 Hz stimulus was delivered for 5 s while images are acquired at 2 Hz in order to select only active synapses. A differential image was constructed by subtracting a five-frame average obtained immediately before the test train of stimulation from a five-frame average obtained just after stimulation. This difference image highlighting the stimulus-dependent increase of fluorescence was subjected to segmentation based on wavelet transform. All identified masks and calculated time courses were visually inspected for correspondence to individual functional pre-synaptic boutons. The mask was then transferred to the images acquired every 3 ms during a 1 Hz electrical stimulation. Successful fusion events were those where the fluorescence intensity of the first point following stimulation was greater than thrice the standard deviation of 200 points prior the increase in fluorescence. The measurement of release probability was made according to the number of successful responses over the total number of stimulations applied.

#### Modeling

Computer modeling was performed using the MCell/CellBlender simulation environment (http://mcell.org) with MCell version 3.3, CellBlender version 1.1, and Blender version 2.77a (http://blender.org). The realistic model of glutamatergic synaptic environment was constructed from 3D-EM of hippocampal area CA1 neuropil as described in 37–39. The 3D-EM reconstruction contains all plasma membrane bounded components including dendrites, axons, astrocytic glia and the extracellular space itself, in a 6×6×5 um3 volume of hippocampal area CA1 stratum radiatum from adult rat. The AMPAR chemical kinetic properties were obtained from the well-established model published in Jonas et al., 1993, and the kinetic parameters were adjusted to fit with the recorded mEPSCs (see 9). Three surface properties are defined. The synapse, the PSD (identified on EM data) and a 100 nm domain inside the PSD which correspond to the AMPAR nanodomain. According to literature and to our dSTORM data, 200 PSD-95 molecules were released. They freely diffuse inside the PSD, and are palmitoylated at a certain rate (kon=35, koff=0.7) when they enter inside the nanodomain area. These kinetic rate constants result in a steady-state accumulation of around 70 palmitoylated PSD-95 inside the nanodomain according to experimental results. PSD-95 can be also inactivated with a certain rate to mimic LTD. Concerning AMPAR, a total of 120 receptors were released at time zero and were distributed in two separate pools as follow: one pool of 60 AMPARs were allowed to diffuse on the membrane surface and a second pool of 60 AMPARs represented the endocytosed state. Individual receptors exchange between the pools at equal forward and backward rates to maintain equilibrium. To mimic LTD induction, the rate of exchange to the endocytosed state was increased. At the surface, AMPARs are mobile with a diffusion coefficient of 0.5 µm^2^.s-1, and can interact inside the nanodomain with palmitoylated PSD-95 (kon = 5, koff = 1). This interaction slows down the AMPAR to 0.005 µm^2^.s-1 and slows down the AMPAR diffusion constant to retain the molecular complex within the nanodomain leading to an equilibrium of around 20 to 25 AMPAR trapped into the domain (similar to results described in the literature). All this organization of PSD-95 and AMPAR at the synapse were simulated at a time step of 1 ms for 50 000 iterations (50 ms), until reaching an equilibrium (as illustrated figure 6S1). Then the simulations were switched to a time step of 1 µs for 250 000 iterations to model the AMPAR responses when the glutamate is released at the pre-synaptic level, in front of the nanodomain. We tested 3 individual modifications of the kinetic rate constants to mimic LTD. (1) A multiplication by 3 of the endocytosis rate mimics the endocytosis-dependent LTD (ATP type or initial phase of NMDA type of LTD). (2) A 4 times increase of the PSD-95 + AMPAR koff, which favors the untrapping of AMPAR. (3) A 4-time increase of the inactivation rate of PSD-95, mimicking the decrease of PSD-95 experimentally observed. 100 simulation trials are averaged for each conditions, and 2500 glutamate are releases for each release event.

#### Sampling and Statistics

Summary statistics are presented as mean ± SEM (Standard Error of the Mean). Statistical significance tests were performed using GraphPad Prism software (San Diego, CA). Normality tests were performed with D’Agostino and Pearson omnibus tests. For non-normally distributed data, we applied Mann-Whitney test or Wilcoxon matched-pairs signed rank tests for paired observations. When the data followed normal distribution, we used paired or unpaired t-test for paired observations unless stated otherwise. ANOVA test was used to compare means of several groups of normally distributed variables. Indications of significance correspond to p values <0.05(*), p < 0.005(**), and p<0.0005(***). After ANOVA analysis, we apply a Dunnett’s post test to determine the p value between two conditions, results of these tests are noted Anova post-test.

