## Supplemental Figures for "Specific nanoscale synaptic reshuffling and control of short-term plasticity following NMDAR- and P2XR-dependent Long-Term Depression"

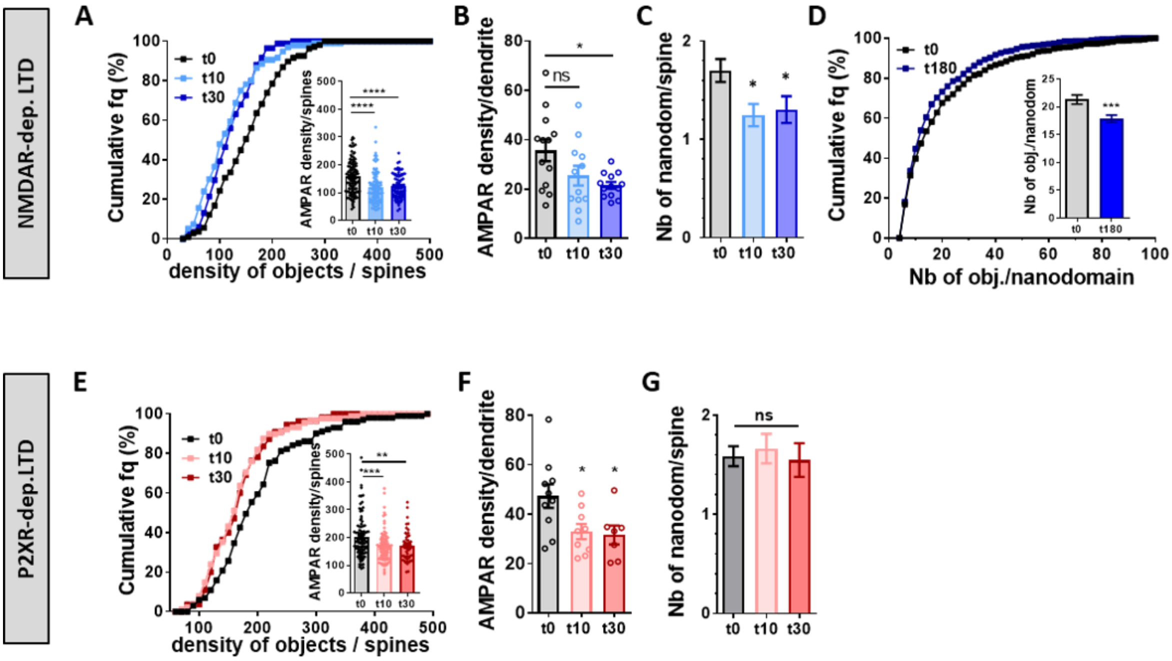


**Figure 1 Suppl.1. NMDA and ATP application triggers a rapid and long lasting decrease of AMPAR surface expression and nanoscale organization.**

(A) Cumulative distribution of AMPAR density per dendritic spine (n=107, 96 and 84 for t0, t10 and t30 respectively), and in the inset, the average histogram. The density of AMPARs per spine was measured 0, 10 and 30 minutes following NMDA treatment (mean +/- SEM, one-way ANOVA, $p<0.0001$ and Dunnett's post-test found significant differences between t0 and t10 or t30, $p<0.0001$). AMPAR surface expression is significantly decreased in dendritic spines 10 and 30 minutes following NMDA treatment compared to non-treated cells. (B) Average of AMPAR density per dendrite measured 0, 10 and 30 minutes following NMDA treatment (mean +/- SEM, n=13, 12 and 13 respectively, one-way ANOVA, p=0.0183 and Dunnett's post-test found significant difference between t0 and t30 p=0.0118 but not between t0 and t10, p=0.0851). AMPAR surface expression is significantly decreased in neuronal dendritic shafts 30 minutes following NMDA treatment compared to non-treated cells. (C) Average of AMPAR nanodomain number per dendritic spine measured 0, 10 and 30 minutes following NMDA treatment (mean +/- SEM, n=107, 105 and 79 respectively, one-way ANOVA, p=0.0123 and Dunnett's post-test found significant difference between t0 and t10 or t30, p=0.0112 and p=0.0465 respectively). AMPAR nanodomain number per dendritic spine is significantly decreased in neuronal dendritic shafts 10 and 30 minutes following NMDA treatment compared to non-treated cells. (D) Cumulative distribution of nanodomain AMPAR content (n=556 and 544 for t0 and t180 respectively), and in the inset, the average histogram. The number of AMPARs per nanodomains was measured at basal state (t0) and 180 minutes following NMDA treatment (mean +/- SEM, n=556 and 544, unpaired t-test, p=0.0010). Nanodomain content is significantly decreased 180 minutes following NMDA treatment compared to non-treated cells.

(E-G) Similar experiments as from A to C has been realized using ATP treatment to trigger LTD. (E) Cumulative distribution of AMPAR density per dendritic spine (n=101, 88 and 55 for t0, t10 and t30 respectively), and in the inset, the average histogram. The density of AMPARs per spines was measured 0, 10 and 30 minutes following ATP treatment (mean +/- SEM, one-way ANOVA, p=0.0005 and Dunnett's post-test found significant differences between t0 and t10 or t30, p=0.0008 and p=0.0052 respectively). AMPAR surface expression is significantly decreased in dendritic spines 10 and 30 minutes following ATP treatment compared to non-treated cells. (F) Average of AMPAR density per dendrite measured 0, 10 and 30 minutes following ATP treatment (mean +/- SEM, n=10, 9 and 7 respectively, one-way ANOVA, p=0.0186 and Dunnett's post-test found significant difference between t0 and t10 or t30 p=0.0304 and p=0.0266 respectively). AMPAR surface expression is significantly decreased in neuronal dendritic shafts 10 and 30 minutes following ATP treatment compared to non-treated cells. (G) Average of AMPAR nanodomain number per dendritic spine measured 0, 10 and 30 minutes following ATP treatment (mean +/- SEM, n=82, 68 and 55 respectively, one-way ANOVA, p=0.8399). AMPAR nanodomain number per dendritic spine is maintained 10 and 30 minutes following ATP treatment compared to non-treated cells.

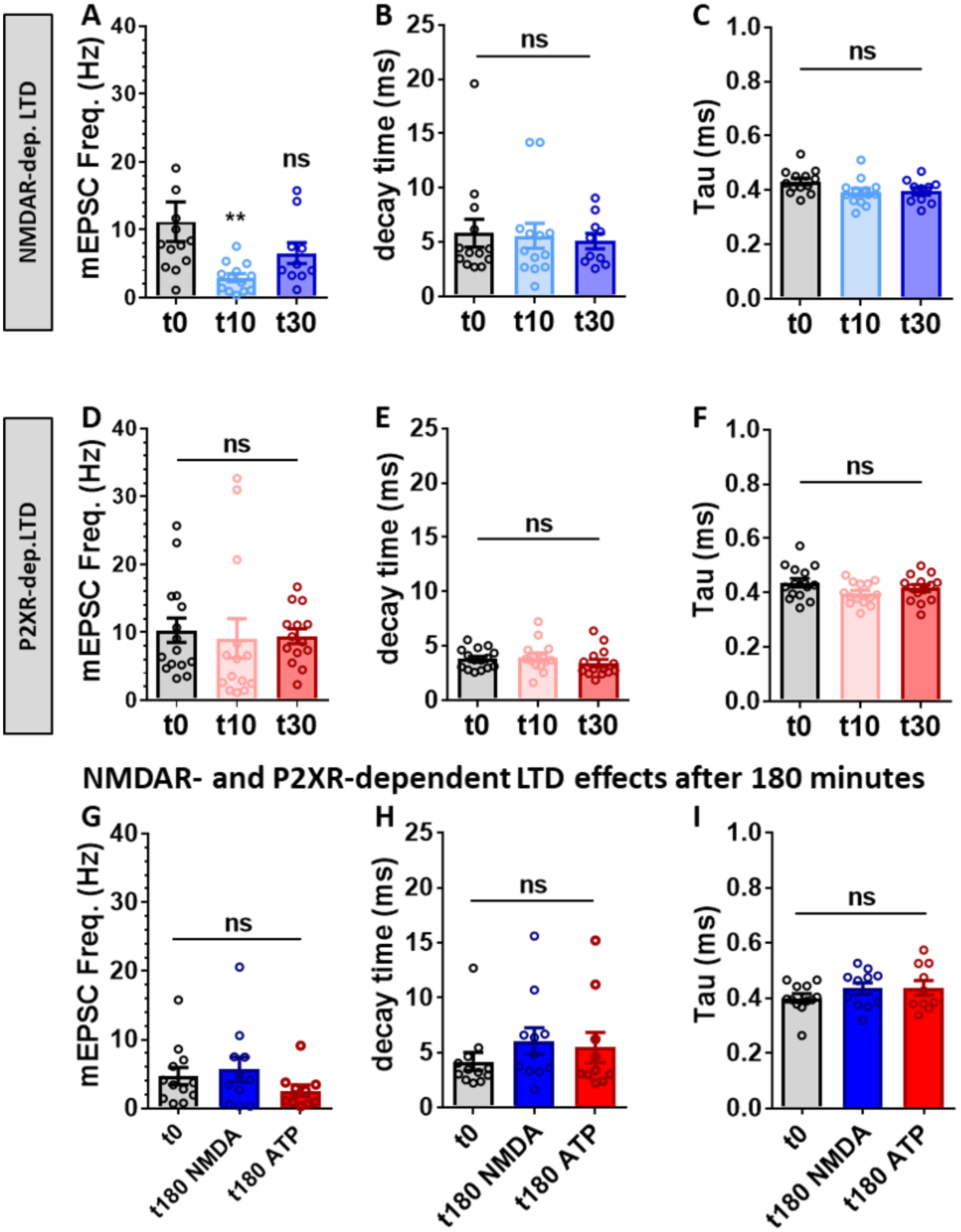


**Figure 1 Suppl.2. Impact of NMDA and ATP treatment on mEPSC properties.**

(A-C) mEPSC frequency (A), decay time (B) and tau (C) at basal state (t0) or 10 and 30 minutes after NMDA treatment (n=13, 13 and 10 respectively, (A) one-way ANOVA, p=0.0179 with Dunnett's post-test showing a significant difference between t0 and t10, p=0.0094; (B,C) one-way ANOVA, p=0.9033 and p=0.0707 respectively. (D-F) mEPSC frequency (D), decay time (E) and tau (F) at basal state (t0) or 10 and 30 minutes after ATP treatment (n=15, 14 and 14 respectively, one-way ANOVA, p=0.0905, p=0.4583 and p=0.1274 respectively). (G-I) mEPSC frequency (G), decay time (H) and tau (I) at basal state (t0) or 180 minutes after NMDA (t180 NMDA) or ATP (t180 ATP) treatments (n=12, 11 and 10 respectively, one-way ANOVA, p=0.2920, p=0.4848 and p=0.3639 respectively).

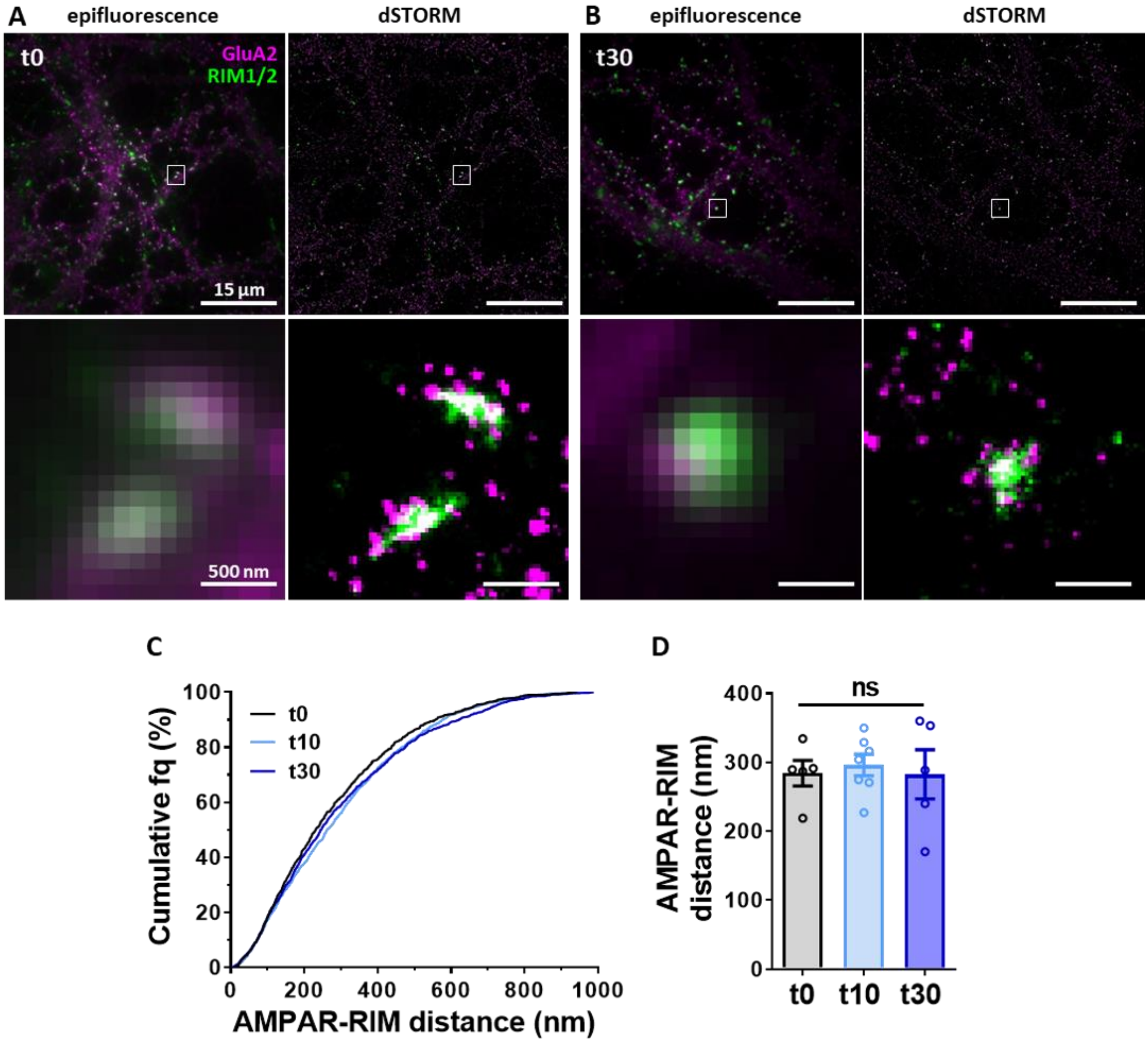


**Figure 1 Suppl.3. NMDA treatment does not impact on the colocalization of the pre-synaptic RIM and the post-synaptic AMPAR domains**

(A and B) Example of dual color D-STORM images with RIM1/2 in green and GluA2-containing AMPARs in purple before treatment (A) and 30 minutes after NMDA application (B). Top Right panels are low resolution images, top left panels are dSTORM reconstructed images, bottom are zoom on synapses from both low and high resolution images. Cumulative distribution (C) and Average per cell (D) of the AMPAR-RIM cluster distances (centroid to centroid) at t0 and 10 and 30 minutes following NMDA treatment. No significant differences are measured (one-way ANOVA, p=0.897).

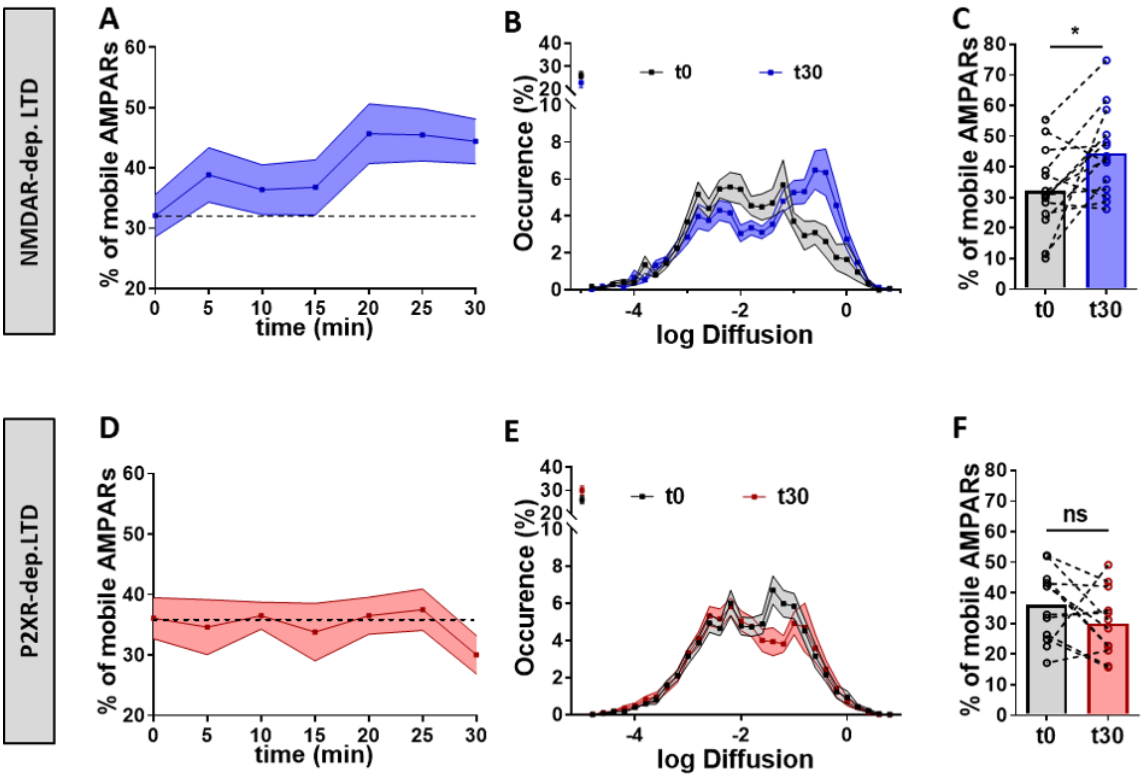


**Figure 2 Suppl.1. NMDAR-dependent LTD but not P2XR-dependent LTD is associated to a long term increase of synaptic AMPAR lateral diffusion.**

(A) Time-lapse (from 0 to 30 minutes) of synaptic GluA2-containing AMPAR mobility following NMDAR-dependent LTD induction protocol (blue line) (n=14). (B) Average distribution of the log(D), (D being the diffusion coefficient of endogenous AMPAR synaptic trajectories) in control condition (black line) and 30 minutes after NMDA treatment (blue line). (C) Average of the mobile fraction at synapses per cell, before and 30 minutes after NMDA treatment (n=14 cells, mean +/- SEM, paired t-test, p=0.0115). (D) Time-lapse (from 0 to 30 minutes) of synaptic GluA2-containing AMPAR mobility following P2XR-dependent LTD induction (red line) (n=14). (E) Average distribution of the log(D) in control condition (black line) and 30 minutes after ATP treatment (red line). (F) Average of the mobile fraction at synapses per cell, before and 30 minutes after ATP treatment (n=14 cells, mean +/- SEM, paired t-test, p=0.1563).

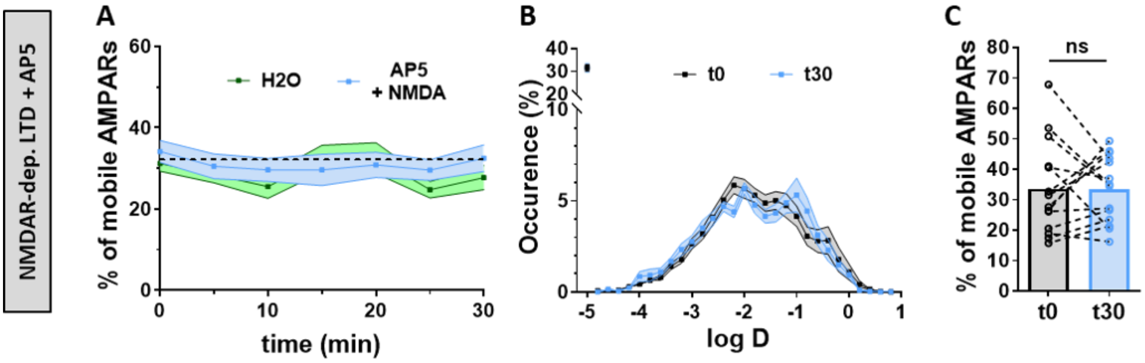


**Figure 2 Suppl.2. AMPAR increased mobility after NMDA treatment is dependent of NMDAR specific activation.**

(A) Time-lapse (from 0 to 30 minutes) of GluA2-containing AMPAR mobility following NMDAR-dependent LTD induction in the presence of AP5 (a specific NMDAR antagonist, light blue line) compared to vehicle application (green line) (n=12 and 10 respectively). No change in GluA2-containing AMPAR mobility occurs after NMDA application in the presence of AP5 (50 µM). (B) Average distribution of the log(D) in control condition (black line) and 30 minutes after NMDA treatment in presence of AP5 (light blue line). (C) Average of the mobile fraction per cell, before and 30 minutes after NMDA treatment with AP5 (n=14 cells, mean +/- SEM, paired t-test, p=0.9697).

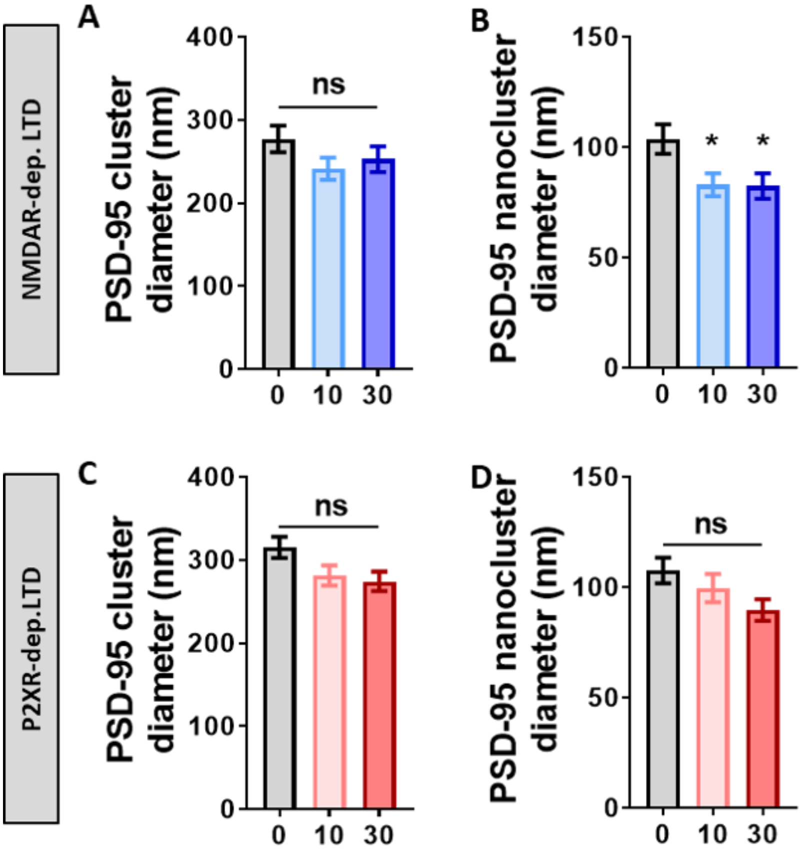


**Figure 3.Suppl.1. Impact of NMDA - or ATP treatment on PSD-95 cluster and nanocluster properties.**
(A and B) Average size of PSD-95 molecules per cluster (A) and per nanoclusters (B) in basal state, 10 and 30 minutes after NMDA treatment (mean +/- SEM, n=17, 19 and 18 respectively, one-way ANOVA p=0.2338 for clusters; one-way ANOVA p=0.0212 and Dunnett's post-test found significant between t0 and t10 and between t0 and t30 conditions, p=0.0297 and p=0.0268 respectively, for nanoclusters). (C and D) Average size of PSD-95 molecules per cluster (C) and per nanoclusters (D) in basal state, 10 and 30 minutes after ATP treatment (mean +/- SEM, n=18, 19 and 16 respectively, one-way ANOVA p=0.0504 for clusters; one-way ANOVA p=0.1165 for nanoclusters).

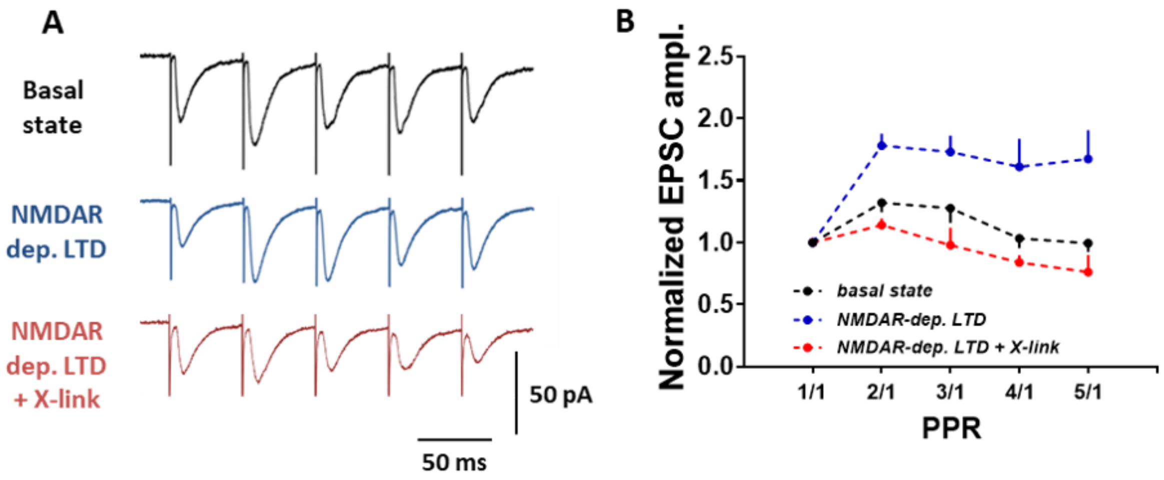


**Figure 4.Suppl.1. AMPAR crosslinking blocks the frequency stimulation facilitation associated to NMDA-dependent LTD.**

(A) Average of the 5 EPSC amplitudes, normalized by the first response intensity. Paired-pulse stimulation was performed on acute hippocampal slices either untreated (basal state), or 30 minutes after NMDAR-dependent LTD induction in presence of anti-GFP antibody (control) or anti-GluA2 antibody (0.3µg/µL, inducing AMPAR cross-link). Injection of antibodies have been done in the stratum radiatum area of the whole-cell patch neuron 20 minutes after the NMDAR-dependent LTD induction. As described in Heine et al. 2008 and Constals et al. 2015, GluA2 cross-linking decrease the PPR.

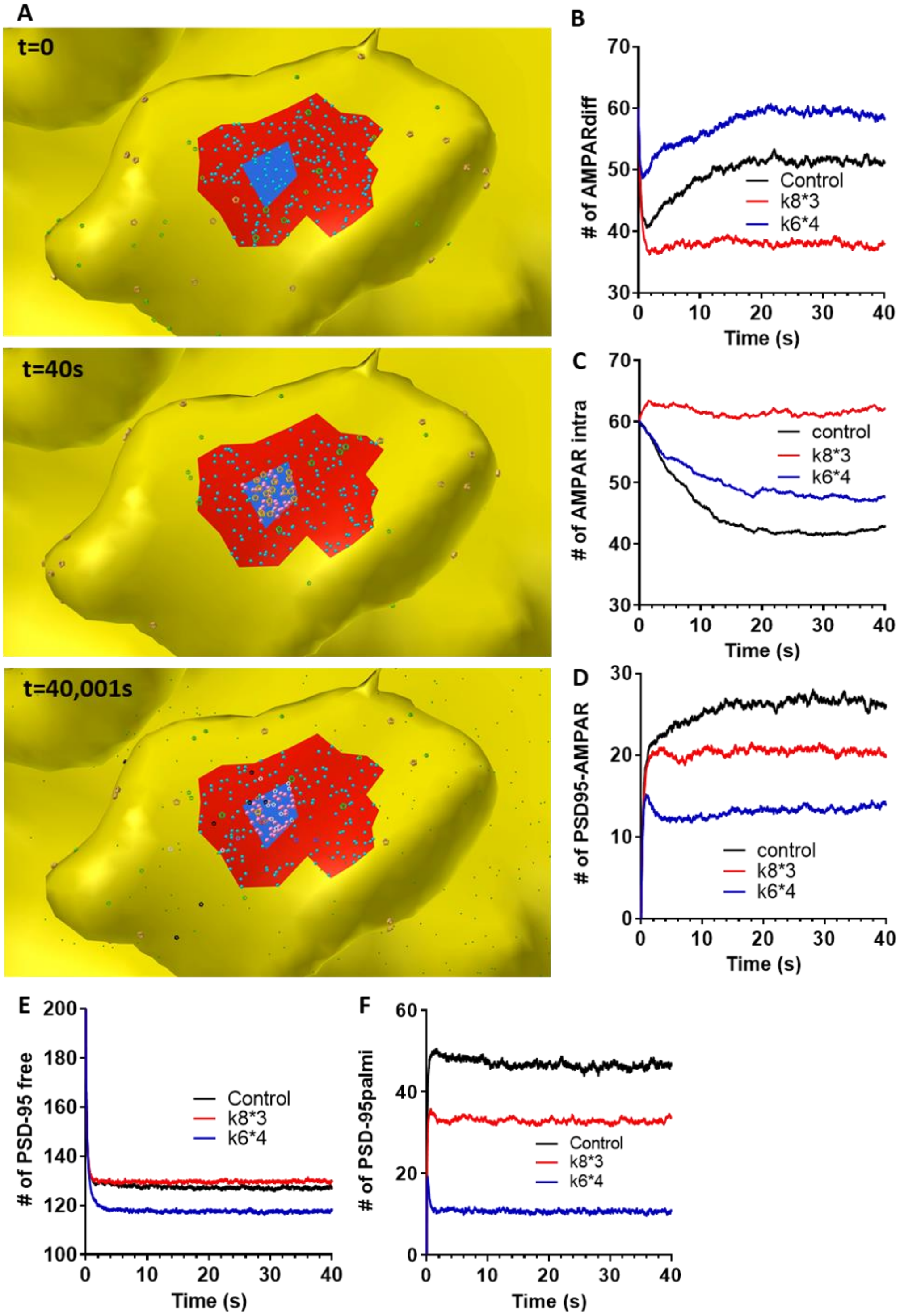


**Figure 5.Suppl.1. Effect of in silico LTD on the equilibrium between various state of both PSD-95 and AMPAR** (A) Example of images obtained with the model at t=0 (top panel), t=40 s when protein organization reach a stable state (middle panel) and at t=40.001 s, 1 ms after first glutamate release (bottom panel). Icosahedrons represent AMPAR, and colors differ in function of their states: orange for the closed, green for the endocytosed, white for the opened, black for the desensitized, etc. Dots represent PSD-95, blue for the freely diffusive and pink for the palmitoylated. (B-F) Kinetics of accumulation of the various protein species in control or when LTD is mimicked by either an increase of endocytosis (red line, k8*3) or by an inactivation of PSD-95 (blue line, k6*4). We report, the evolution of the number of: diffusive AMPAR (B), internalized AMPAR (C), PSD-95 coupled to AMPAR (D), free PSD-95 (E) and palmitoylated PSD-95 (F). The proportion of each species at the equilibrium are closed to the values experimentally obtained.

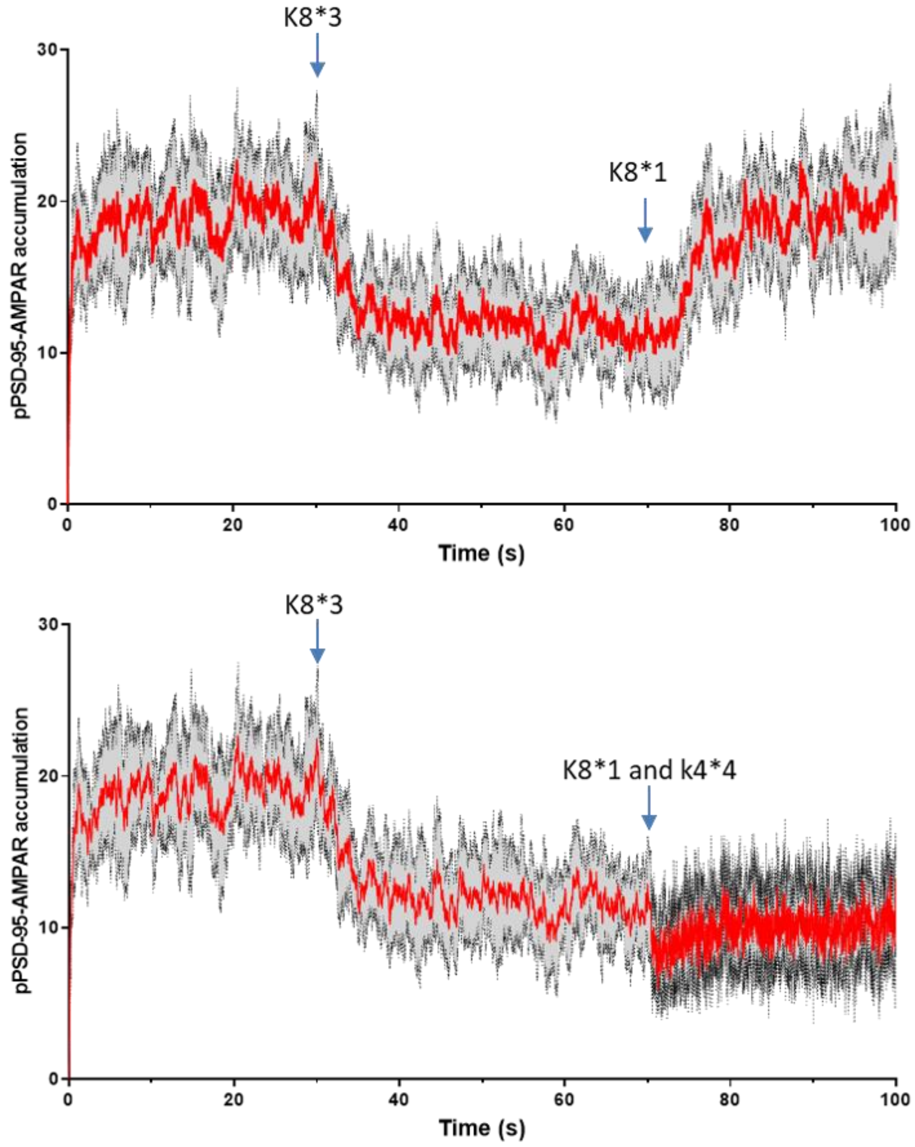


**Figure 5.Suppl.2. Simulation shows that LTD-induced increase of endocytosis need to be maintained or compensate to stabilize depression of AMPAR**

(A) Simulation of AMPAR accumulation in nanodomain. For the first 30 s we observe the recruitment of AMPAR. At 30 s, the endocytosis rate is multiply by 3 to mimic an LTD. At 70 s the endocytosis rate is returned to its initial value, triggering to a progressive replenishment of the AMPAR nanodomain. (B) Similar modeling are realized but at 70 s, we re-initiate endocytosis rate and in parallel we decreased the affinity of AMPAR for the traps (as shown Figure 6C). We observed a stabilization of the nanodomain depletion, and interestingly, to an increase of the noise due to the more rapid exchange of AMPAR.
